# Manifold Learning for fMRI time-varying FC

**DOI:** 10.1101/2023.01.14.523992

**Authors:** Javier Gonzalez-Castillo, Isabel Fernandez, Ka Chun Lam, Daniel A Handwerker, Francisco Pereira, Peter A Bandettini

## Abstract

Whole-brain functional connectivity (*FC*) measured with functional MRI (fMRI) evolve over time in meaningful ways at temporal scales going from years (e.g., development) to seconds (e.g., within-scan time-varying *FC* (*tvFC*)). Yet, our ability to explore *tvFC* is severely constrained by its large dimensionality (several thousands). To overcome this difficulty, researchers seek to generate low dimensional representations (e.g., *2D* and *3D* scatter plots) expected to retain its most informative aspects (e.g., relationships to behavior, disease progression). Limited prior empirical work suggests that manifold learning techniques (*MLTs*)—namely those seeking to infer a low dimensional non-linear surface (i.e., the manifold) where most of the data lies—are good candidates for accomplishing this task. Here we explore this possibility in detail. First, we discuss why one should expect tv*FC* data to lie on a low dimensional manifold. Second, we estimate what is the intrinsic dimension (i.e., minimum number of latent dimensions; *ID*) of *tvFC* data manifolds. Third, we describe the inner workings of three state-of-the-art *MLTs*: Laplacian Eigenmaps (*LE*), T-distributed Stochastic Neighbor Embedding (*T-SNE*), and Uniform Manifold Approximation and Projection (*UMAP*). For each method, we empirically evaluate its ability to generate neuro-biologically meaningful representations of *tvFC* data, as well as their robustness against hyper-parameter selection. Our results show that *tvFC* data has an *ID* that ranges between 4 and 26, and that *ID* varies significantly between rest and task states. We also show how all three methods can effectively capture subject identity and task being performed: *UMAP* and *T-SNE* can capture these two levels of detail concurrently, but L*E* could only capture one at a time. We observed substantial variability in embedding quality across *MLTs*, and within-*MLT* as a function of hyper-parameter selection. To help alleviate this issue, we provide heuristics that can inform future studies. Finally, we also demonstrate the importance of feature normalization when combining data across subjects and the role that temporal autocorrelation plays in the application of *MLTs* to *tvFC* data. Overall, we conclude that while *MLTs* can be useful to generate summary views of labeled *tvFC* data, their application to unlabeled data such as resting-state remains challenging.

## Introduction

From a data-science perspective, an fMRI scan is a four-dimensional tensor with the first three dimensions encoding position in space (*x,y,z*) and the fourth dimension referring to time (*t*). Yet, for operational purposes, it is often reasonable to merge the three spatial dimensions into one and conceptualize this data as a matrix of space vs. time. With current technology (e.g., voxel size ∼ 2×2×2mm^3^, TR ∼ 1.5 s), a representative ten-minutes fMRI scan with full brain coverage will generate a matrix with approximately 400 temporal samples (number of acquisitions) in each of over 40,000 grey matter (GM) locations (number of voxels). Before this data is ready for interpretation, it must undergo several transformations that address three key needs: 1) removal of signal variance unrelated to neuronal activity; 2) spatial normalization into a common space to enable comparisons across subjects and studies; and 3) generation of intuitive visualizations for explorative or reporting purposes. These three needs and how to address them will depend on the nature of the study (e.g., bandpass filtering may be appropriate for resting-state data and not task, the *MNI152* template (Fonov et al., 2009) will be a good common space to report adult data yet not for a study conducted on a pediatric population). This work focuses on how to address the third need—the generation of interpretable visualizations—particularly for studies that explore the temporal dynamics of the human functional connectome.

Most human functional connectome studies use the concept of a functional connectivity matrix (*FC* matrix) or its graph equivalent. Two of the most common types of *FC* matrices in the fMRI literature are: 1) static *FC* (*sFC*) matrices designed to capture average levels of inter-regional activity synchronization across the duration of an entire scan, and 2) time-varying FC (*tvFC*) matrices meant to retain temporal information about how connectivity strength fluctuates as scanning progresses. These two matrix types not only differ on their representational goal, but also in their structure and dimensionality. In a *sFC* matrix, rows (*i*) and columns (*j*) represent spatial locations (e.g., voxels, regions of interest (ROIs)), and the value of a given cell *(i,j)* is a measure of similarity (e.g., Pearson’s correlation, partial correlation, mutual information) between the complete time series recorded at these two locations. When *FC* is expressed in terms of Pearson’s correlation (the most common approach in the fMRI literature), *sFC* matrices are symmetric with a unit diagonal. Moreover, they can be transformed from their original *2D* form (*N x N*; *N* = number of spatial locations) into a *1D* vector of dimensionality:

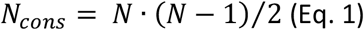

with *N*_*cons*_ being the number of unique pair-wise connections.

Conversely, a *tvFC* matrix is a much larger data structure where rows represent connections, and columns represent time (Fig. 1.A). The size of a *tvFC* (*N*_*cons*_ *x N*_*wins*_) is determined as follows. The number of rows (*N*_*cons*_) is given by the number of spatial locations contributing to the matrix (Eq. 1). The number of columns (*N*_*wins*_) is a function of the duration of the scan (*N*_*acq*_ = number of temporal samples or acquisitions) and the mechanism used to construct the matrix, which in fMRI, is often some form of sliding window technique that proceeds as follows. First, a temporal window of duration shorter than the scan is chosen (e.g., *W*_*duration*_ = 20 samples << *N*_*acq*_). Second, a *sFC* matrix is computed using only the data within that temporal window. The resulting *sFC* matrix is then transformed into its *1D* vector representation, which becomes the first column of the *tvFC* matrix. Next, the temporal window slides forward a given amount determined by the windowing step (e.g., *W*_*step*_ = 3 samples), a new *sFC* matrix is computed for the new window, transformed into its *1D* form, and inserted as the second column of the *tvFC* matrix. This process is repeated until a full window can no longer be fit to the data. This results in *N*_*wins*_ columns, with *N*_*wins*_ given by:

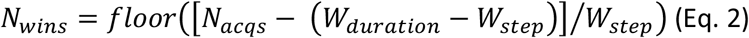

Figure 1.A shows a *tvFC* matrix for a 25-minute-long fMRI scan (*N*_*acqs*_=1017, TR=1.5s) acquired continuously as a subject performed and transitioned between four different tasks (i.e., rest, working memory, arithmetic calculations, and visual attention; (Gonzalez-Castillo et al., 2015)). Each task was performed continuously for two separate three minutes periods. The *tvFC* matrix was generated using a brain parcellation with 157 ROIs and a sliding window approach (*W*_*duration*_ = 30 samples, *W*_*step*_ = 1 sample). As such, the dimensions of the *tvFC* matrix are *12246 connections X 988 temporal windows*. Figure 1.A shows the matrix at scale (each datapoint represented by a square) so we can appreciate the disproportionate ratio between number of connections (Y-axis) and number of temporal samples (X-axis). Figure 1.B shows the same matrix as Fig. 1.A, but this time the temporal axis has been stretched so that we can better observe the temporal evolution of *FC*. In this view of the data, connections are sorted according to average strength across time. The colored segments on top of the matrix indicate the task being performed at a given window (gray=rest, blue=working memory [WM], yellow=visual attention [VA], green=arithmetic [Math]). Colors are only shown for task-homogenous windows, meaning those that span scan periods when the subject was always performing the same task (i.e., no transitions or two different tasks). Transition windows, namely those that include more than one task and/or instruction periods, are marked as empty spaces between the colored boxes. Figure 1.C and 1.D show the same *tvFC* matrix as Fig. 1.A, but with connections sorted by temporal volatility (i.e., coefficient of variation) and hemispheric membership respectively. In all instances, irrespective of sorting, it is quite difficult to directly infer from these matrices basic characteristics of how *FC* varies over time and/or relates to behavior. This is in large part due to the high dimensionality of the data.

**Figure 1.**
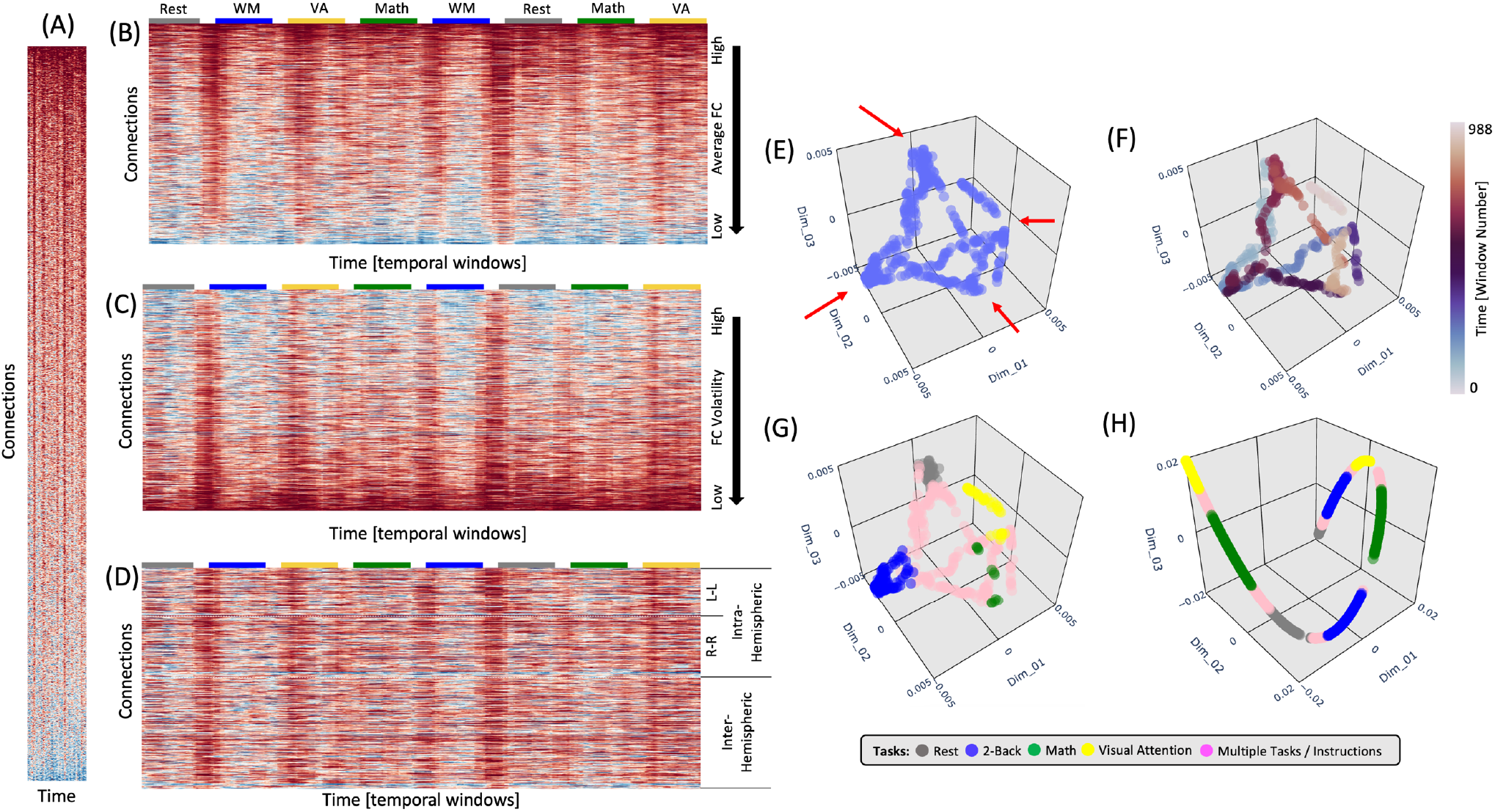
(A) tvFC matrix at scale to illustrate the disproportionate larger dimensionality of the connectivity axis (y-axis) relative to the time axis (x-axis). (B) Same *tvFC* matrix as in (A) but no longer at scale. The x-axis has not been stretched to better observe how connectivity evolves over time. Connections are sorted in terms of their average strength. Task-homogenous windows are clearly marked above the *tvFC* matrix with color-coded rectangles. (C) Same *tvFC* matrix with connections sorted in terms of their volatility (as indexes by the coefficient of variance). (D) Same *tvFC* matrix with connections sorted according to hemispheric membership. Intra-hemispheric connections appear at the top of the matrix and inter-hemispheric at the bottom. (E,F,G) Laplacian Eigenmap (correlation distance, k=90) for the *tvFC* matrix in (A-D) with no color annotation (E), annotated by time (F), and annotated by task (G). (H) Laplacian Eigenmap for the *tvFC* matrix in (A-D) using correlation distance and k=20.

When an initial exploration of high dimensional data is needed, it is common practice to generate a low dimensional representation (e.g., two or three dimensions) that can be easily visualized yet preserves important information about the structure of the data (e.g., groups of similar samples, presence of outliers) in the original space. Figure 1.E shows one such representation of our *tvFC* matrix generated using a manifold learning method called *Laplacian Eigenmaps* (*LE*; (Belkin and Niyogi, 2003)). In this representation, each column from the *tvFC* matrix becomes a point in *3D* space. In other words, each point represents the brain *FC* during a portion of the scan (in this case a 30 samples window). Points that are closer in this lower dimensional space are supposed to correspond to whole brain *FC* patterns that are similar. A first look at Figure 1.E reveals that there are four different recurrent *FC* configurations (corners marked with red arrows). Moreover, if we annotate points with colors that represent time (Figure 1.F), we can also learn that each of those configurations were visited twice, once during the first half of the scan (blue tones) and a second time during the second half (red tones). Finally, if we annotate the points with information about the task being performed at each window, we can clearly observe that the four corners correspond to *FC* patterns associated with each of the tasks—with temporally separated occurrences of the task appearing close to each other—and that transitional windows tend to form trajectories going between two corners corresponding to the tasks at both ends of each transitional period.

To achieve the meaningful representations presented in Figure 1.E-G, one ought to make several analytical decisions beyond those related to fMRI data preprocessing and brain parcellation selection. These include the selection of a dimensionality reduction method, a dissimilarity function and a set of additional method-specific hyper-parameters (e.g., number of neighbors, perplexity, learning rate, etc.). In the same way to how using the wrong bandpass filter while pre-processing resting-state data can eliminate all neuronally-relevant information, choosing incorrect hyper-parameters for *LE* can produce less meaningful low dimensional representations of *tvFC* matrices. Figure 1.H shows one such instance, where using an excessively small neighborhood (*K*_*nn*_) resulted on a *3D* scatter that only captures temporal autocorrelation and misses the other important data characteristics discussed in the previous paragraph (e.g., the repetitive task structure of the experimental paradigm).

In this manuscript we will explore the usability of three prominent manifold learning methods—namely *LE*, T-distributed Stochastic Neighbor Embedding ((Maaten and Hinton, 2008); *T-SNE)*, and Uniform Manifold Approximation and Projection ((McInnes et al., 2018); *UMAP)*—to generate low-dimensional representations of *tvFC* matrices that retain neurobiological and behavioral information. These three methods were selected because of their success across scientific disciplines, including many biomedical applications (Diaz-Papkovich et al., 2019; Kollmorgen et al., 2020; Zeisel et al., 2018). First, we will introduce the manifold hypothesis and the concept of intrinsic dimension (*ID*) of a dataset. We will also provide a programmatic level description (as opposed to purely mathematical) of each of these methods. Next, we will evaluate each method’s ability to generate meaningful low dimensional representations of *tvFC* matrices using a clustering framework and a predictive framework for both individual scans and the complete dataset. Our results demonstrate the *tvFC* data reside in low dimensional manifolds that can be effectively estimated by the three methods under evaluation, yet also highlight the importance of correctly choosing key hyper-parameters as well as that of considering the effects of temporal autocorrelation when designing experiments and interpreting the final embeddings. In this regard, we provide a set of heuristics that can guide their application in future studies. In parallel, we also demonstrate the value of *ID* for deciding how many dimensions ought to be explored or included in additional analytical steps (e.g., spatial transformations, classification), and demonstrate its value as an index of how *tvFC* complexity varies between resting and task states.

## Theory

### Manifold Hypothesis

The manifold hypothesis sustains that many high dimensional datasets that occur in the real world (e.g., real images, speech, etc.) lie along or near a low-dimensional manifold (e.g., the equivalent of a curve or surface beyond three dimensions) embedded in the original high dimensional space (often referred to as the ambient space). This is because the generative process for real world data usually has a limited number of degrees of freedom constrained by laws (e.g., physical, biological, linguistic, etc.) specific to the process. For example, images of human faces lie along a low dimensional manifold within the higher dimensional ambient pixel space because most human faces have a quite regular structure (one nose between two eyes sitting above a mouth, etc.) and symmetry. This makes the space of pixel combinations that lead to images of human faces a very limited space compared to that of all possible pixel combinations. Similarly, speech lies in a low dimensional manifold within the higher dimensional ambient space of sound pressure timeseries because speech sounds are restricted both by the physical laws that limit the type of sounds the human vocal tract can generate and by the phonetic principles of a given language. Now, does an equivalent argument apply to the generation of *tvFC* data? In other words, is there evidence to presume that *tvFC* data lies along or near a low dimensional manifold embedded within the high dimensional ambient space of all possible pair-wise *FC* configurations? The answer is yes. Given our current understanding of the functional connectome and the laws governing fMRI signals it is reasonable to assume that the manifold hypothesis applies to fMRI-based *tvFC* data.

First, the topological structure of the human functional connectome is not random but falls within a small subset of possible topological configurations known as small-world networks, which are characterized by high clustering and short path lengths (Sporns and Honey, 2006). This type of network structure allows the co-existence of functionally segregated modules (e.g., visual cortex, auditory cortex) yet also provide efficient ways for their integration when needed. Second, *FC* is constrained by anatomical connectivity (Goñi et al., 2014); which is also highly organized and far from random. Third, when the brain engages in different cognitive functions, *FC* changes accordingly (Gonzalez-Castillo and Bandettini, 2018); yet those changes are limited (Cole et al., 2014; Krienen et al., 2014), and global properties of the functional connectome, such as its small-worldness, are preserved as the brain transitions between task and rest states (Bassett et al., 2006). Forth, *tvFC* matrices have structure in both the connectivity and time dimensions. On the connectivity axis, pair-wise connections tend to organize into networks (i.e., sets of regions with higher connectivity among themselves than to the rest of the brain) that are reproducible across scans and across participants. On the temporal axis, connectivity time-series show temporal autocorrelation due to the sluggishness of the hemodynamic response and the use of overlapping sliding windows. Fifth, previous attempts at applying manifold learning to *tvFC* data have proven successful at generating meaningful low dimensional representations that capture differences in *FC* across mental states (Bahrami et al., 2019; Gao et al., 2021; Gonzalez-Castillo et al., 2019), sleep stages (Rué-Queralt et al., 2021), and populations (Bahrami et al., 2019).

### Intrinsic Dimension

One important property of data is their intrinsic dimension (*ID*, (Campadelli et al., 2015)), namely the minimum number of variables (i.e., dimensions) required to describe the manifold where the data lie with little loss of information. Given the above-mentioned constrains that apply to the generative process of *tvFC* data, it is reasonable to expect that the intrinsic dimension of *tvFC* data will be significantly smaller than that of the original ambient space, yet it may still be a number well above three. Because *ID* informs us about the minimum number of variables needed to faithfully represent the data, having an estimate of what is the *ID* of *tvFC* data is key. For example, if the *ID* is greater than three, one should not restrict visual exploration of low dimensional representations of *tvFC* data to the first three dimensions and should also explore dimensions above those. Similarly, if manifold estimation is used to compress the data or extract features for a subsequent classification step, knowing the *ID* can help us decide how many dimensions (i.e., features) to keep. Finally, *ID* of a dataset can also be thought of as a measure of the complexity of the data (Ansuini et al., 2019). In the context of *tvFC* data, such a metric might have clinical and behavioral relevance.

Intrinsic dimension estimation is currently an intense area of research (Albergante et al., 2019; Facco et al., 2017), with *ID* estimation methods in continuous evolution to address open issues such as computational complexity, under-sampled distributions, and dealing with datasets that reside in multiple manifolds (see (Campadelli et al., 2015) for an in-depth review of these issues). Because no consensus exists on how to optimally select an *ID* estimator, here we will compare estimates from three state-of-the-art methods, namely local PCA (*lPCA;* (Fan et al., 2010)), two nearest neighbors (*twoNN;* (Facco et al., 2017)) and Fisher Separability (*FisherS;* (Albergante et al., 2019)). These three *ID* estimators were selected because of their complementary nature on how they estimate *ID*, robustness against data redundancy and overall performance (Bac et al., 2021). In all instances, we will report both global and local *ID* estimates. The global *ID* (*ID*_*global*_) is a single *ID* estimate per dataset generated using all data samples. It works under the assumption that the whole dataset has the same *ID* (see Supplementary Figure 1.C for a counter example). Conversely, local *ID* (*ID*_*local*_) estimates are computed on a sample-by-sample basis using small vicinities of size determined by the number of neighbors (*k*_*nn*_). In that way, local *ID* estimates can help identify regions with different *IDs*; yet its accuracy is more dependent on noise levels and the relative size of the data curvature with respect to *k*_*nn*_. Please see Supplementary Note 1 for additional details on the relationship between *ID*_*global*_ and *ID*_*local*_, and about how *k*_*nn*_ can affect *ID*_*local*_ estimation.

### Laplacian Eigenmaps

The first method that we evaluate is the Laplacian Eigenmap (*LE*) algorithm originally described by Belkin et al. (2003) and publicly available as part of the *scikit-learn* library (Pedregosa et al., 2011). In contrast to linear dimensionality reduction methods (e.g., *PCA*) that seek to preserve the global structure of the data, *LE* attempts to preserve its local structure. Importantly, this bias towards preservation of local over global structure facilitates the discovery of natural clusters in the data.

The *LE* algorithm starts by constructing an undirected graph (*G*) from the data (Fig. 2.D). In *G*, each node represents a sample (i.e., a column of the *tvFC* matrix; Fig 2.A), and edges are drawn only between nodes associated with samples that are “*close*” in original space. The construction of *G* proceeds in two steps. First, a dissimilarity matrix (*DS*) is computed (Fig. 2.B). For this, one must choose a distance function (*d*). Common choices include the *Euclidean, Correlation*, and *Cosine* distances (see Suppl. Note 2 for additional details about these distance metrics). Next, this *DS* matrix is transformed into an affinity matrix (*W*) using the *N-nearest neighbors* algorithm (Fig. 2.C). In *W*, the entry for *i* and *j* (*W*_*ij*_) is equal to 1 (signaling the presence of an edge) if and only if node *i* is among the *K*_*nn*_ nearest neighbors of node *j* (*i* → *j*) or *j* is among the *K*_*nn*_ nearest neighbors of node *i (j* → *i)*. Otherwise *W*_*ij*_ equals zero. An affinity matrix constructed this way is equivalent to an undirected, unweighted graph (Fig. 2.D). According to Belkin et al., it is also possible to work with a weighted version of the graph, for example:

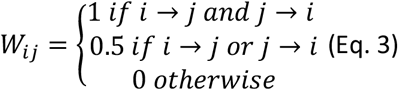

This alternative version is the one used in the implementation of the *scikit-learn* library used in this work.

**Figure 2.**
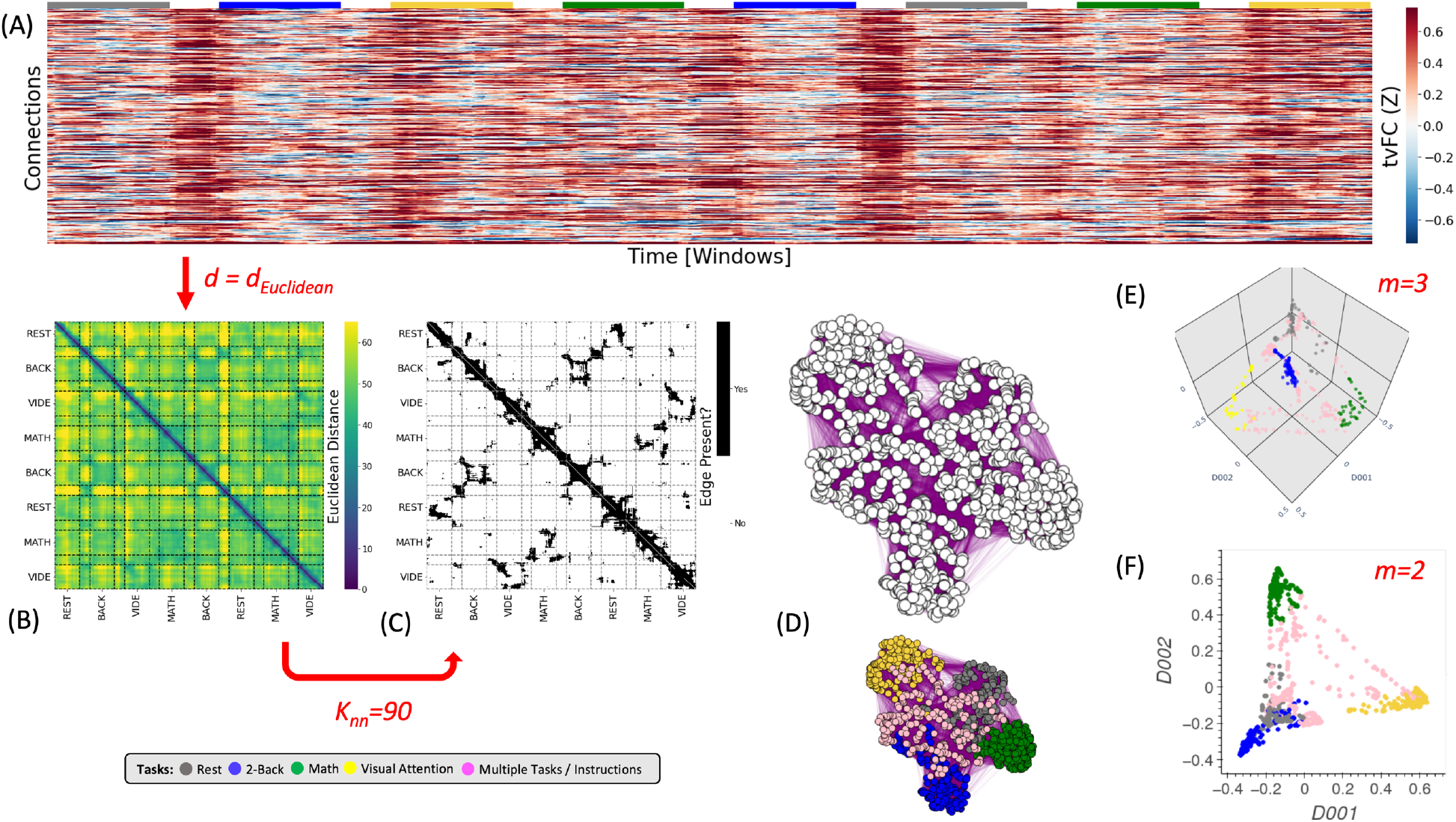
The Laplacian Eigenmap Algorithm. (A) Representative tvFC matrix for a multi-task run. Columns indicate windows and rows indicate connections. (B) Dissimilarity matrix for the tvFC matrix in (A) computed using the Euclidean distance function. (C) Affinity matrix computed for (B) using *K*_*nn*_=90 neighbors. Black cells indicate 1 (i.e., an edge exists) and white indicate zero (no edge). (D) Graph visualization of the affinity matrix in (C). In the top graph all nodes are colored in white to highlight that the *LE* algorithm used no information about the tasks at any moment. The bottom graph is the same as the one above, but now nodes are colored by task to make apparent how the graph captures the structure of the data (e.g., clusters together windows that correspond to the same experimental task). (E) Final embedding for m = 3. This embedding faithfully presents the task structure of the data. (F) Final embedding for m = 2. In this case, the windows for rest and memory overlap. Red arrows and text indicate decision points in the algorithm. Step-by-step code describing the *LE* algorithm and used to create the different panels of this figure can be found in the code repository that accompanies this publication (*Notebook N02_Figure02_Theory_LE.ipynb*)

Once the graph is built, the next step is to obtain the Laplacian matrix (*L*)^1^ of the graph, which is defined as *L* = *D* − *W* (Eq. 4). In eq. 4, *W* is the affinity matrix (Eq. 3), and *D* is a matrix that holds information about the degree (i.e., number of connections) of each node on the diagonal and zeros elsewhere. The last step of the *LE* algorithm is to extract eigenvalues (*λ*_0_ ≤ *λ*_1_ ≤ ⋯ ≤ *λ*_*k*−1_) and eigenvectors (*f*_0_, …, *f*_*k*−1_) by solving *Lλ* = *λDf* (Eq. 5). Once those are available, the embedding of a given sample *x* in a lower dimensional space with *m* ≪ *k* dimensions is given by:

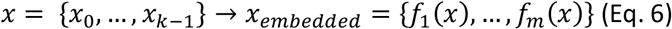

The first eigenvector *f*_0_ is ignored because its associated eigenvalue *λ*_0_ is always zero. For those interested in a step-by-step mathematical justification of why the spectral decomposition of *L* renders a representation of the data that preserves local information, please read *Section 3* of the original work by Belkin et al. Intuitively, the *Laplacian* matrix is a linear operator that holds information about all between-sample relationships in the manifold and the eigenvectors obtained via its spectral decomposition provide a set of orthonormal bases.

In summary, the *LE* algorithm requires, at a minimum, the selection of a distance function (*d*), and a neighborhood size (*k*_*nn*_). These two hyper-parameters (marked in red in Fig. 2) determine the construction of *G* because they mathematically specify what it means for two *tvFC* patterns to be similar (or graph nodes to be connected). Because the *LE* algorithm does not look back at the input data once *G* is constructed, and all algorithmic steps past the construction of *G* are fixed, appropriately selecting *d* and *k*_*nn*_ is key for the generation of biologically meaningful embeddings of *tvFC* data. Finally, as with any dimensionality reduction technique, one must also select how many dimensions to explore (*m;* Fig. 2.E-F), but in the case of *LE* such decision does not affect the inner workings of the algorithm.

### T-Stochastic Nearest Neighbor Embedding (T-SNE)

The second technique evaluated here is *T-SNE* (Maaten and Hinton, 2008), which is a commonly used method for visualizing high dimensional biomedical data in two or three dimensions. Like *LE, T-SNE*’s goal is to generate representations that give priority to the preservation of local structure. These two methods are also similar in that their initial step requires the selection of a distance function used to construct a dissimilarity matrix (*DS; Fig. 3.B*) that will subsequently be transformed into an affinity matrix (*P; Fig. 3.C*). Yet, *T-SNE* uses a very different approach to go from *DS* to *P*. Instead of relying on the *N*-nearest *neighbor* algorithm, *T-SNE* models pair-wise similarities in terms of probability densities. Namely, the affinity between two points *x*_*i*_ and *x*_*j*_ in original space is given by the following conditional *Gaussian* distribution:

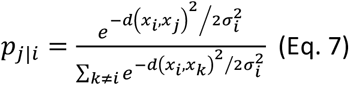

As such, *T-SNE* conceptually defines the affinity between *x*_*i*_ and *x*_*j*_ as the likelihood that *x*_*i*_ would pick *x*_*j*_ as its neighbor, with the definition of neighborhood given by a *Gaussian* kernel of width 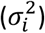 centered at *x*_*i*_. The width of the kernel 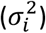 is sample-dependent to accommodate datasets with varying densities (see Suppl. Fig. 2 for an example) and ensure all neighborhoods are equivalent in terms of how many samples they encompass. In fact, *T-SNE* users do not set 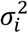 directly but select neighborhood size via the *perplexity (PP)* parameter, which can be thought of as an equivalent to *k*_*nn*_ in the *LE* algorithm. Please see Supplementary Note 3 for more details on how perplexity relates to neighborhood size.

**Figure 3.**
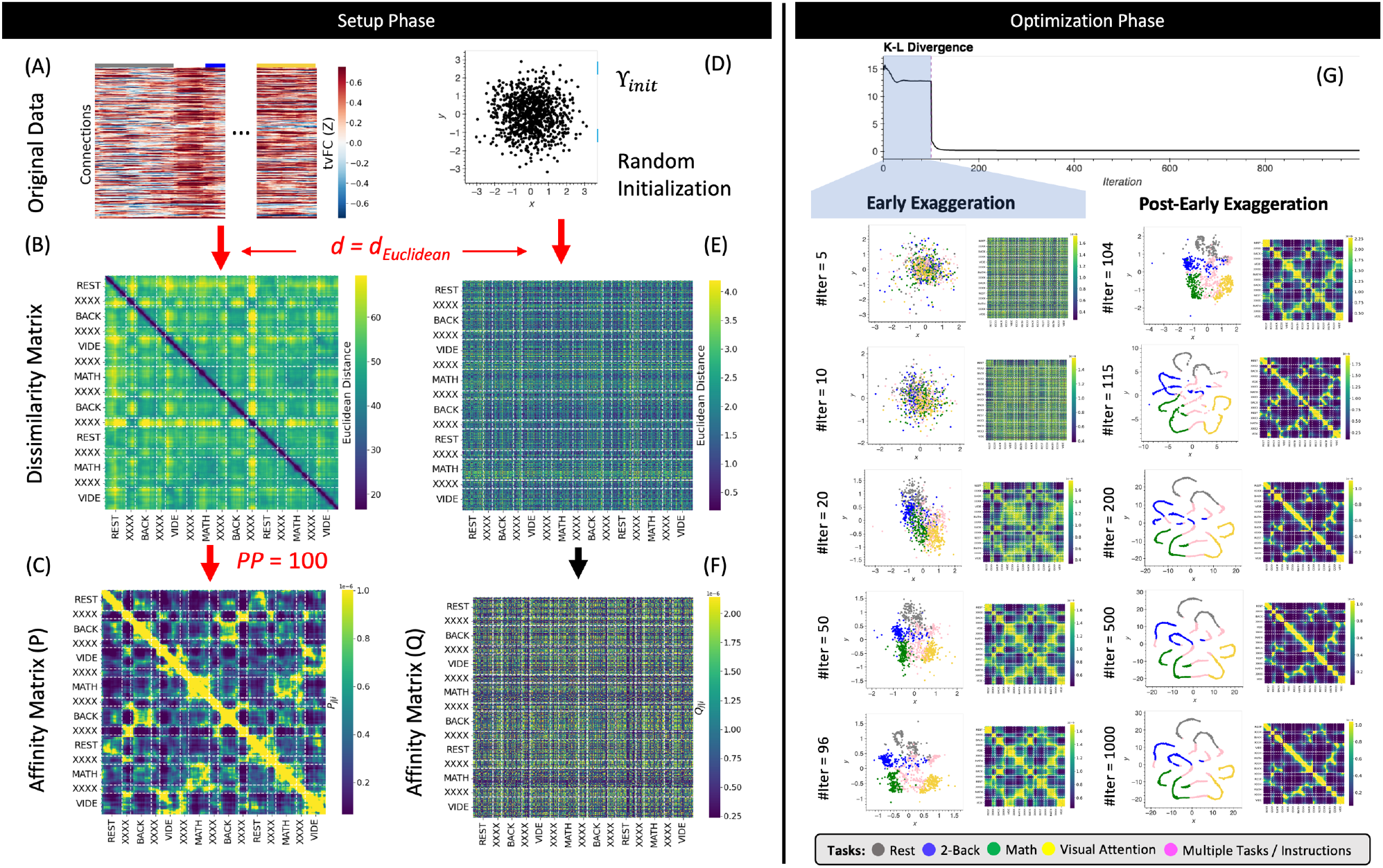
The *T-SNE* Algorithm. (A) Representative *tvFC* matrix for a multi-task run. Columns indicate windows and rows indicate connections. (B) Dissimilarity matrix for the *tvFC* matrix in (A) computed using the Euclidean distance function. (C) Affinity matrix generated using equations 3 and 4 and a perplexity value of 100. (D) Random lower-dimensional representation of the data. Each dot represents one column of the data in (A). (E) Dissimilarity matrix for the data in (D) computed using the *Euclidean* distance function. (F) Affinity matrix for the data in (D) computed using a T-distribution function as in equation 5. (G) Evolution of the cost function with the number of gradient descent iterations. In this execution, the early exaggeration factor was set to 4 for the initial 100 iterations (as originally described by (2008)). A dashed vertical line marks the iteration when the early exaggeration factor was removed (early exaggeration phase highlighted in light blue). Below the cost function evolution curve, we show embeddings and affinity matrices for a set of representative iterations. Iterations corresponding to the early exaggeration periods are shown on the left, while iterations for the post early exaggeration period are shown on the right. In the embeddings, points are colored according to the task being performed on each window. Windows that contain more than one task are marked in pink with the label “XXXX”. Step-by-step code describing a basic implementation of the *T-SNE* algorithm and used to create the different panels of this figure can be found in the code repository that accompanies this publication (*Notebook N03_Figure03_Theory_TSNE.ipynb*)

Because conditional probabilities are not symmetric, entries in the affinity matrix *P* are defined as follows in terms of the conditional probabilities defined in Eq. 7:

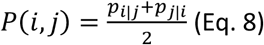

*T-SNE* proceeds next by generating an initial set of random coordinates for all samples in the lower *m* dimensional space (γ_*init*_; Fig. 3.D). Once γ_*init*_ is available, a dissimilarity (Fig 3.E) and an affinity matrix (*Q;* Fig 3.F) are also computed for this random initial low dimensional representation of the data. To transform *DS* into *Q, T-SNE* uses a *T-Student* distribution (Eq. 9) instead of a *Gaussian* distribution. The reason for this is that *T-Student* distributions have heavier tails than *Gaussian* distributions, which in the context of *T-SNE*, translates into higher affinities for distant points. This gives *T-SNE* the ability to place distant points further apart in the lower dimensional representation and use more space to model the local structure of the data.

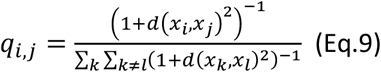

The *T-SNE* steps presented so far constitute the setup phase of the algorithm (Fig 3.A-F). From this point onward (Fig 3.G), the *T-SNE* algorithm proceeds as an optimization problem where the goal is to update γ (e.g., the locations of the points in lower *m* dimensional space) in a way that makes *P* and *Q* most similar (e.g., match pair-wise distances). *T-SNE* solves this optimization problem using gradient descent to minimize the *Kullback*-Leibler (*KL*) divergence between both affinity matrices (Eq. 10).

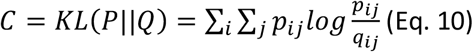

Figure 3.G shows how *C* evolves with the number of gradient descent iterations. Below the *C* curve we show intermediate low dimensional representations (γ) and affinity matrices (*Q*) at representative iterations. In this execution of *T-SNE*, it takes approx. 20 to 50 iterations for γ to present some meaningful structure: rest windows (grey dots) being together on the top left corner of the embedding and visual attention windows (yellow dots) being on the bottom right corner. As the number of iterations grows the profile becomes more distinct, and windows associated with the math (green) and working memory tasks (blue) also start to separate. Because *T-SNE*’s optimization problem is non-convex, modifications to the basic gradient descent algorithm are needed to increase the chances of obtaining meaningful low dimensional representations. These include, among others, early compression (i.e., the addition of an L2-penalty term to the cost function during initial iterations), early exaggeration (i.e., a multiplicative term on *P* during initial iterations), and an adaptative learning rate procedure. For example, figure 3.G shows the effects of early exaggeration. At iteration 100, when early exaggeration is removed, *C* sharply decreases as *P* suddenly becomes much closer in value to *Q*. As the optimization procedures continues beyond that point, we can observe how the temporal autocorrelation inherent to *tvFC* data dominates the structure of the embedding (e.g., continuous spaghetti-like structure), yet windows corresponding to the two different periods of each task still appear close to each other showing the ability of the embedding to also preserve behaviorally relevant information.

These optimization “*tricks*” result into additional hyper-parameters that can affect the behavior of *T-SNE*, yet not all of them are always accessible to the user. For example, in the *scikit-learn* library (the one used in this work), one can set the early exaggeration factor, but not the number of gradient descent iterations to which it applies. Given optimization of gradient descent is beyond the scope of this work, here we focus our investigation only on the effects of distance metric (*d*), perplexity (*PP*), and learning rate (β). Similarly, it is also worth noting that the description of *T-SNE* provided in this section corresponds roughly to that originally proposed by Van der Maaten and Hinton (2008). Since then, several modifications have been proposed, such as the use of the *Burnes-Hut* approximation to reduce computational complexity (Maaten, 2014), and the use of *PCA* initialization to introduce information about the global structure of the data during the initialization process (Kobak and Berens, 2019). These two modifications are available in *scikit-learn* and will be used as defaults in the analyses described below.

In summary, although *T-SNE* and *LE* share the goal of generating low dimensional representations that preserve local structure and rely on topological representations (i.e., graphs) to accomplish that goal, the two methods differ in important aspects. First, the number of hyper-parameters is much higher for *T-SNE* than *LE*. This is because the *T-SNE* algorithm contains a highly parametrized optimization problem with options for the learning rate, early exaggeration, early compression, Barnes-Hut radius, and initialization method to use. Second, in *LE* the number of desired dimensions (*m*) is used to select the number of eigenvectors to explore, and as such, it does not affect the outcome of the algorithm in any other way. That is not the case for *T-SNE*, where *m* determines the dimensionality of γ at all iterations, and therefore the space that the optimization problem can explore in search of a solution. In other words, if one were to run LE for *m = 3* and later decide to only explore the first two dimensions, that would be a valid approach as the first two dimensions of the solution for *m = 3* are equivalent to the two dimensions of the solution for *m = 2*. The same is not true for *T-SNE*, which would require separate executions for each scenario (*m = 2* and *m = 3*).

### Uniform Manifold Approximation and Projection (UMAP)

The last method considered here is *UMAP* (McInnes et al., 2018). *UMAP* is, as of this writing, the latest addition to the family of dimensionality reduction techniques based on neighboring graphs. Here we will describe *UMAP* only from a computational perspective that allow us to gain an intuitive understanding of the technique and its key hyperparameters. Yet, it is worth noting that *UMAP* builds on strong mathematical foundations from the field of topological data analysis (*TDA;* (Carlsson, 2009)), and that, it is those foundations (as described in Section 2 of (McInnes et al., 2018)), that justify each procedural step of the *UMAP* algorithm. For those interested in gaining a basic understanding of *TDA* we recommend these two works: (Chazal and Michel, 2021) which is written for data scientists as the target audience, and (Sizemore et al., 2019) which is more specific to applications in neuroscience.

*UMAP* can be described as having two phases: 1) the construction of an undirected weighted graph for the data, and 2) the computation of a low dimensional layout for the graph. Phase one proceeds as follows. First, a dissimilarity matrix (*DS;* Figure 4.B) is computed using the user-selected distance function *d*. This *DS* matrix is then transformed into a binary adjacency matrix (*A;* Figure 4.C) using the *k-nearest neighbor* algorithm and a user-selected number of neighbors (*k*_*nn*_). This is similar to the first steps in *LE*, except that here matrix *A* defines a directed (as opposed to undirected in *LE*) unweighted graph *G*_*a*_ = (*V, E, w*_*a*_) (Figure 4.D) where *V* is a set of nodes/vertices representing each data sample (i.e., a window of connectivity), *E* is the set of edges signaling detected neighboring relationships, and *w*_*a*_ equals 1 for all edges in *E*. Third, *UMAP* computes an undirected weighted graph *G*_*b*_ = (*V, E, w*_*b*_) (Figure 4.F) with the same set of nodes and edges of *G*_*a*_, but with weights *w*_*b*_ given by

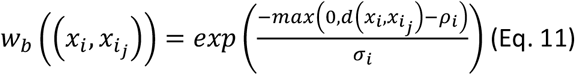

where 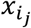refers to the *j*^*th*^ nearest neighbor of node *x*_*i*_ with *j* = {1.. *k*_*nn*_}, 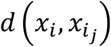 is their dissimilarity as provided by the distance function *d*, and *ρ*_*i*_ and *σ*_*i*_ are node-specific normalization factors given by equations 12 and 13 below.

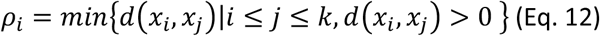

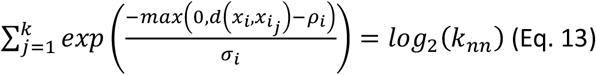

By constructing *G*_*b*_ this way, *UMAP* ensures that in *G*_*b*_ all nodes are connected to at least one other node with a weight of one, and that the data is represented as if it were uniformly distributed on the manifold in ambient space. Practically speaking, equations 11 through 13 transform original dissimilarities between neighboring nodes into exponentially decaying curves in the range [0,1] (Figure 4.E).

**Figure 4.**
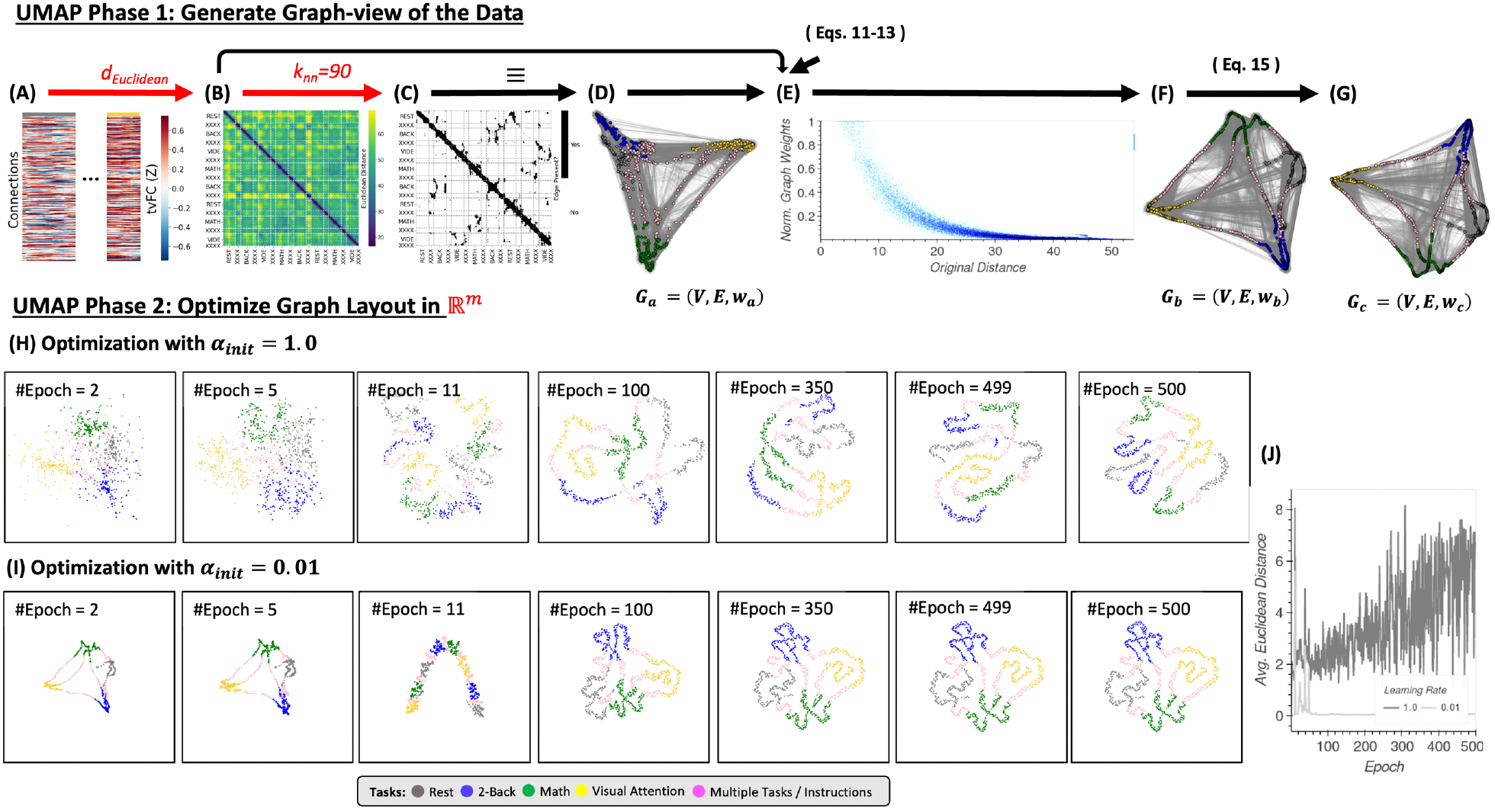
(A) Input tvFC data. (B) Dissimilarity matrix obtained using the Euclidean distance as a distance function. (C) Binary non-symmetric affinity matrix for *K*_*nn*_=90. (D) Graph equivalent of the affinity matrix in (C). (E) Effect of the distance normalization step on the original dissimilarities between neighboring nodes. (F) Graph associated with the normalized affinity matrix. (G) Final undirected graph after application of equation 15. This is the input to the optimization phase. (H) Representative embeddings at different epochs of the stochastic gradient descent algorithm for an initial learning rate equal to 1. (I) Same as (H) but when the initial learning rate is set to 0.01. (J) Difference between embeddings at consecutive epochs measured as the average Euclidean distance across all node locations for an initial learning rate of 1 (dark grey) and 0.01 (light grey). Step-by-step code describing a basic implementation of the T-SNE algorithm and used to create the different panels of this figure can be found in the code repository that accompanies this publication (*Notebook N05_Figure04_Thoery_UMAP.ipynb*)

Finally, if we describe *G*_*b*_ in terms of an affinity matrix *B*

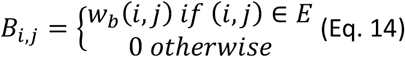

Then, we can generate a symmetrized version of matrix *B*, called *C*, as follows

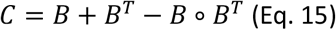

This matrix *C* represents graph *G*_*c*_ = (*V, E, w*_*c*_) (Figure 4.G), which is the undirected weighted graph whose layout is optimized during phase 2 of the UMAP algorithm as described next.

Once *G*_*c*_ is available, the second phase of the *UMAP* algorithm is concerned with finding a set of positions for all nodes ({*Y*_*i*_}_*i*=1..*N*_) in ℝ^*m*^, with *m* being the desired number of user-selected final dimensions. For this, *UMAP* uses a force-directed graph-layout algorithm that positions nodes using a set of attractive and repulsive forces proportional to the graph weights. An equivalent way to think about this second phase of *UMAP*—that renders a more direct comparison with *T-SNE*—is in terms of an optimization problem attempting to minimize the edge-wise cross-entropy (Eq. 16) between *G*_*c*_ and an equivalent weighted graph *H* = (*V, E, w*_*h*_) with node layout given by {*Y*_*i*_}_*i*=1..*N*_ ∈ ℝ^*m*^. The goal being to find a layout for *H* that makes *H* and *G*_*c*_ are as similar as possible as dictated by the edge-wise cross-entropy function.

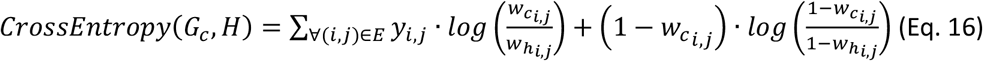

If we compare T-SNE’s (Eq. 10) and UMAP’s (Eq. 16) optimization cost functions, we can see that Eq. 10 is equivalent to the left term of Eq. 16. This left term represents the set of attractive forces along the edges that is responsible for positioning together similar nodes in ℝ^*m*^. Conversely, the right term of Eq. 16 represents a set of repulsive forces between nodes that are responsible for enforcing gaps between dissimilar nodes. This additional term helps UMAP preserve some of the global structure of the data while still capturing local structure.

Figures 4.H&I exemplify how UMAP phase 2 proceeds for two different learning rates (a key hyper-parameter of the optimization algorithm). These two panels in figure 4 show embeddings at representative epochs (e.g., 2,5, 11, 100, 350, 499 and 500). Additionally, figure 4.J shows the average *Euclidean* distance between all nodes at two consecutive epochs. This allows us to evaluate if optimization proceeds smoothly with small changes in the embedding from one step to the next, or abruptly. An abrupt optimization process is not desirable because, if so, a small change in the number of epochs to run can lead to substantially different results. Figure 4.I shows how when the learning rate is set to 0.01, optimization proceeds abruptly only at the initial stages (*N*_*epoch*_ < 100) and then stabilizes. In this case a small change in the maximum number of epochs to run will not affect the results. Moreover, the embedding for *N*_*epoch*_ = 500 in figure 4.I has correctly captured the structure of the data (i.e., the tasks). Next, in figure 4.H we observe that the same is not true for a learning rate of 1 (the default value in the *umap-learn* library). In this case, embeddings substantially change from one epoch to the next all the way to *N*_*epoch*_ = 500. These results highlight the strong effects that hyper-parameters associated with optimization phase of *UMAP* can have when working with *tvFC* data.

In summary, although *UMAP* shares many conceptual and practical aspects with *LE* and *T-SNE*, it differs in important ways on the specifics of how a graph is generated from the data and how this graph is translated into a low dimensional embedding. Similarly to *LE* and T-SNE, *UMAP* requires careful selection of distance metric (*d*), neighborhood size (*k*_*nn*_) and the dimensionality of the final space (*m*). In addition, like *T-SNE, UMAP* exposes many additional parameters associated with its optimization phase. Here we have only discussed the learning rate and maximum number of epochs, but many other are available. For a complete list please check the *umap-learn* documentation. For this work, unless expressed otherwise, we will use default values for all other hyper-parameters. Finally, there is one more hyper-parameter specific to *UMAP* that we have not yet discussed called minimum distance (*min_dist*). Its value determines the minimum separation between closest samples in the embedding. In that way, *min_dist* controls how tightly together similar samples appear in the embedding (see Supplementary Note 4 for additional details).

## Methods

### Dataset

This work is conducted using a multi-task dataset previously described in detail in (Gonzalez-Castillo et al., 2019). In summary, it contains data from 22 subjects (13 females; age 27 ± 5 y.o.) who gave informed consent in compliance with a protocol approved by the Institutional Review Board of the National Institute of Mental Health in Bethesda, MD. The data from two subjects were discarded from the analysis due to excessive spatial distortions in the functional time series.

Subjects were scanned continuously for 25 min and 24 s while performing four different tasks: rest with eyes open (*REST*), simple mathematical computations (*MATH*), 2-back working memory (*WM*), and visual attention/recognition (*VA*). Each task occupied two separate 180-s blocks, preceded by a 12s instruction period. Task blocks were arranged so that each task was always preceded and followed by a different task. Additional details can be found on the supplementary materials accompanying Gonzalez-Castillo et al. (2015).

### Data Acquisition

Imaging was performed on a Siemens 7 T MRI scanner equipped with a 32-element receive coil (Nova Medical). Functional runs were obtained using a gradient recalled, single shot, echo planar imaging (gre-EPI) sequence: (TR = 1.5 s; TE = 25 ms; FA = 50°; 36 interleaved slices; slice thickness = 2 mm; in-plane resolution = 2 × 2 mm; GRAPPA = 2). Each multi-task scan consists of 1017 volumes acquired continuously as subjects engage and transition between the different tasks. In addition, high resolution (1 mm^3^) T1-weighted magnetization-prepared rapid gradient-echo and proton density sequences were acquired for presentation and alignment purposes.

### Data Pre-processing

Data preprocessing was conducted with AFNI (Cox, 1996). Preprocessing steps match those described in Gonzalez-Castillo et al. (2015), and include: (i) despiking; (ii) physiological noise correction (in all but four subjects, due to the insufficient quality of physiological recordings for these subjects); (iii) slice time correction; and (iv) head motion correction. In addition, mean, slow signal trends modeled with Legendre polynomials up to seventh order, signal from eroded local white matter, signal from the lateral ventricles (cerebrospinal fluid), motion estimates, and the first derivatives of motion were regressed out in a single regression step to account for potential hardware instabilities and remaining physiological noise (*ANATICOR*; (Jo et al., 2010)). Finally, time series were converted to signal percent change, bandpass filtered [0.03–0.18 Hz], and spatially smoothed (*FWHM* = 4 mm). The cutoff frequency of the high pass filter was chosen to match the inverse of window length (*WL* = 45s); following recommendations from (Leonardi and Ville, 2014).

In addition, spatial transformation matrices to go from *EPI* native space to Montreal Neurological Institute (*MNI*) space were computed for all subjects following procedures previously described in (Gonzalez-Castillo et al., 2013). These matrices were then used to bring publicly available regions of interest (*ROI*) definitions from MNI space into each subject’s EPI native space.

### Brain Parcellation

We used 157 regions of interest from the publicly available version of the 200 ROI version of the Craddock Atlas (Craddock et al., 2012). The missing 43 ROIs were excluded because they did not contain at least 10 voxels in all subjects’ imaging field of view. Those were ROIs primarily located in cerebellar, inferior temporal and orbitofrontal regions.

### Time-varying Functional Connectivity

First, for each scan we obtained representative timeseries for all ROIs as the spatial average across all voxels part of the ROI using AFNI program *3dNetCorr* (Taylor and Saad, 2013). Next, we computed time-varying functional connectivity for all scans separately using 45 seconds (30 samples) long rectangular windows with an overlap of one sample (1.5s) in *Python*. Windowed functional connectivity matrices were converted to *1D* vectors by taking only unique connectivity values above the matrix diagonal. These were subsequently concatenated into scan-wise *tvFC* matrices of size 12246 Connections X 988 Windows.

Time-varying *FC* matrices computed this way are referred through the manuscript as non-normalized or as-is. Additionally, we also computed normalized versions of those *tvFC* matrices in which all rows have been forced to have a mean of zero and a standard deviation of one across the time dimension. We refer to those matrices as normalized or *Z*-*scored* matrices.

### Intrinsic Dimension

Three different *ID* estimators were used here: local PCA (*lPCA*), two nearest neighbors (*twoNN*) and Fisher separability (*FisherS*). For each method, we computed both local ID (*ID*_*local*_) estimates at each *tvFC* window and global ID estimates (*ID*_*global*_) at the scan level. We then report on the distribution of these two quantities across the whole sample.

### Dimensionality Reduction

We computed low dimensional representations of the data at two different scales: scan-level and group-level. Scan-level embeddings were generated separately for each scan providing as input their respective *tvFC* matrices.

To generate group-level embeddings, meaning embeddings that contain all windows from all scans in the dataset, we used two different approaches:

- “*Concatenate + Embed*”: in this case, we first concatenate all scan-level *tvFC* matrices into a single larger matrix for the whole dataset. We then provide this matrix as input to the dimensionality reduction step.
- “*Embed + Procrustes*”: here, we first compute scan-level embeddings separately for each scan and then apply the *Procrustes* transformation to bring all of them into a common space.

Table 1 summarizes the different configurations being explored for each manifold learning method.

**Table 1.** Hyper-parameter exploration space

| Hyper-parameters | LE | T-SNE | UMAP |
| --- | --- | --- | --- |
| **Distance Function** | Euclidean, Correlation, Cosine | Euclidean, Correlation, Cosine | Euclidean, Correlation, Cosine |
| **Neighborhood Size** | $K_{nn}$ : [5..200, step=5] | $PP$: [5..100, step=5] + [150,175,200] | $K_{nn}$ : [5..200, step=5] |
| **# Dimensions** | 2,3,5,10,15,20,25,30 | 2,3,10,15 | 2,3,5,10,15,20,25,30 |
| **Learning Rate** | N/A | 10, 50, 75, 100, 200, 500, 1000 | 0.01, 0.1, 1.0 |
| **Minimum Distance** | N/A | N/A | 0.8 |

### Embedding Evaluation

Two different frameworks—clustering and predictive—were used to quantitatively evaluate the quality of the resulting low dimensional representations. The clustering framework looks at the ability of those representations to show groupings of *FC* configurations that match labels of interest (e.g., task being performed). The use of this framework is primarily motivated by the concept of *FC* states (Allen et al., 2014; Gonzalez-Castillo et al., 2015)—namely short-term recurrent *FC* configurations—and the fact that external cognitive demands modulate *FC* (Gonzalez-Castillo and Bandettini, 2018). As such, a meaningful low dimensional representation of the multi-task dataset should show cluster structure that relates to the different tasks. A common way to measure the cluster consistency in machine learning is the *Silhouette Index* (*SI*; (Rousseeuw, 1987)), which is a measure of cluster cohesion (how similar members of a cluster are to each other) against cluster separation (the minimum distance between samples from two different clusters). *SI* ranges from -1 to 1, with higher *SI* values indicating more clearly delineated clusters. *SI* was computed using the *Python scikit*-*learn* library. Only task-homogenous windows—namely those that do not include instruction periods or more than one task—are used for the computation of the *SI*. For scan-level results we computed *SI* based on tasks. For group-level results we computed *SI* based on tasks and subject identity.

We also evaluate embeddings using a predictive framework. In this case, the goal is to quantify how well low dimensional representations of *tvFC* data performs as inputs to subsequent regression/classification machinery. This framework is motivated by the wide-spread use of *FC* (both static and time-varying) patterns as input features for the prediction of clinical conditions (Rashid et al., 2016), clinical outcomes (Dini et al., 2021), personality traits (Hsu et al., 2018), behavioral performance (Jangraw et al., 2018) and cognitive abilities (Finn et al., 2014). Our quality measure under this second framework is the *F1* accuracy of classification at predicting task name for task-homogenous windows using as input group-level embeddings. We restricted analyses to *UMAP* and *LE* group-level embeddings obtained using the “Embed + *Procrustes”* approach because those have good task separability scores and are computationally efficient at estimating embeddings beyond 3 dimensions. The classification engine used is a logistic regression machine with an *L1* regularization term as implemented in the *scikit-learn* library. We split the data into training and testing sets within a split-half cross-validation procedure. First, we trained the classifier using all windows within the first half of all scans and tested on the remaining of the data. We then switched training and testing sets. Reported accuracy values are the average of the results across the two halves, when they were the test set. It is worth noting that, by splitting the data this way, we achieve two goals: 1) training on data from all tasks in all subjects, and 2) testing using data from windows that are fully temporally independent from the ones used for training.

### Null Models

Two null models were used as control conditions in this study. The first null model (labeled “*randomized connectivity*”) proceeds by randomizing row ordering separately for each column of the *tvFC* matrix. By doing so, the row-to-connection relationship inherent to *tvFC* matrices is destroyed and a given row of the *tvFC* matrix no longer represents the true temporal evolution of *FC* between a given pair of ROIs.

The second model (labeled “*phase randomization*”) proceeds by randomizing the phase of the *ROI* timeseries prior to the computation of the *tvFC* matrix (Handwerker et al., 2012). More specifically, for each ROI we computed its *Fourier* transform, kept the magnitude of the transform but substituted the phase by uniformly distributed phase spectra, and finally applied the inverse *Fourier* transform to get the surrogate ROI representative timeseries. This procedure ensures the surrogate data will retain the autoregressive properties of the original time series, yet the precise timing of signal fluctuations is destroyed.

## Results

### Intrinsic Dimension

Average estimates of global (*ID*_*global*_) and local intrinsic dimension (*ID*_*local*_) across all scans are presented in Figure 5.A-B. We show estimates based on three *ID* estimators: *Local PCA, Two Nearest Neighbors* and *Fisher Separability*. Average *ID*_*global*_ ranged from 26.25 dimensions [*Local PCA*, normalized *tvFC* matrices] to 4.10 dimensions [*Fisher Separability*, no normalization]. *ID*_*global*_ estimates were significantly larger for *Local PC*A than for the two other methods (*p*_*Bonf*_*<1e*^*-4*^). Normalization of *tvFC* matrices had a negligible effect of *ID*_*global*_ estimates. Despite the differences across estimation techniques, in all instances the *ID*_*global*_ of these data is shown to be several orders of magnitude below that of ambient space (i.e., 12246 Connections).

**Figure 5.**
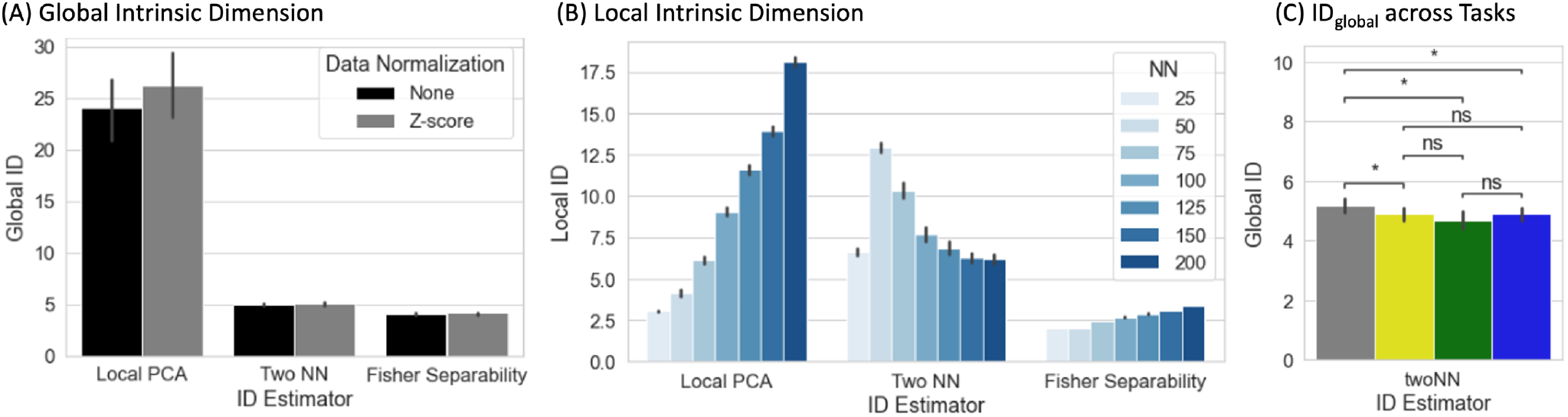
(A) Summary view of local ID estimates segregated by estimation method (Local PCA, Two Nearest Neighbors, Fisher Separability) and normalization approach (None or Z-scoring). Bar height corresponds to average across scans, bars indicate 95% confidence intervals. (B) Summary view of global ID estimates segregated by estimation method and neighborhood size (*NN* = number of neighbors) for the non-normalized data. (C) Statistical differences in global ID across tasks (\**p*_*FDR*_<0.05) [Rest = gray, Visual Attention = yellow, Math = green, 2-Back = blue].

Estimating *ID*_*local*_ requires the selection of a neighborhood size defined in terms of the number of neighbors (*NN*). Figure 5.B shows average *ID*_*local*_ estimates for non-normalized data across all scans as a function of both estimation technique and *NN*. Overall, *ID*_*local*_ ranges between 2 [*Fisher Separability, NN = 50*] and 21 [*Local PCA, NN = 200*]. As it is the case with *ID*_*global*_, *ID*_*local*_ estimates vary significantly between estimators. In general, “*Local PCA”* leads to the largest estimates. Also, there is a general trend for *ID*_*local*_ estimates to increase monotonically with neighborhood size. Exceptions to these two trends only occur for the “*Two NN*” estimator when *NN* ≤ 75. It is important to note that *ID*_*local*_ estimates are always below their counterpart *ID*_*global*_ estimates.

We also computed *ID*_*global*_ separately for each task. When using the *twoNN* estimator, we found that rest has a significantly higher *ID*_*global*_ than all other tasks (Figure 5.C). Such significant difference was not detected with the other two methods.

### Evaluation for Visualization/Exploration Purposes

Figure 6.A-C shows the distribution of *SI*_*task*_ for *2D* and *3D* single-scan embeddings for both original data and the two null models. Each panel shows results for a different *MLT. SI*_*task*_ of original data often reached values above 0.4 (black arrows). That is not the case for either null model. Yet, while the “*Connectivity Randomization*” model always resulted in *SI*_*task*_ near or below zero, the “*Phase Rand*omization” model shows substantial overlap with the lower end of the distribution for original data, especially for *LE* and *UMAP*.

**Figure 6.**
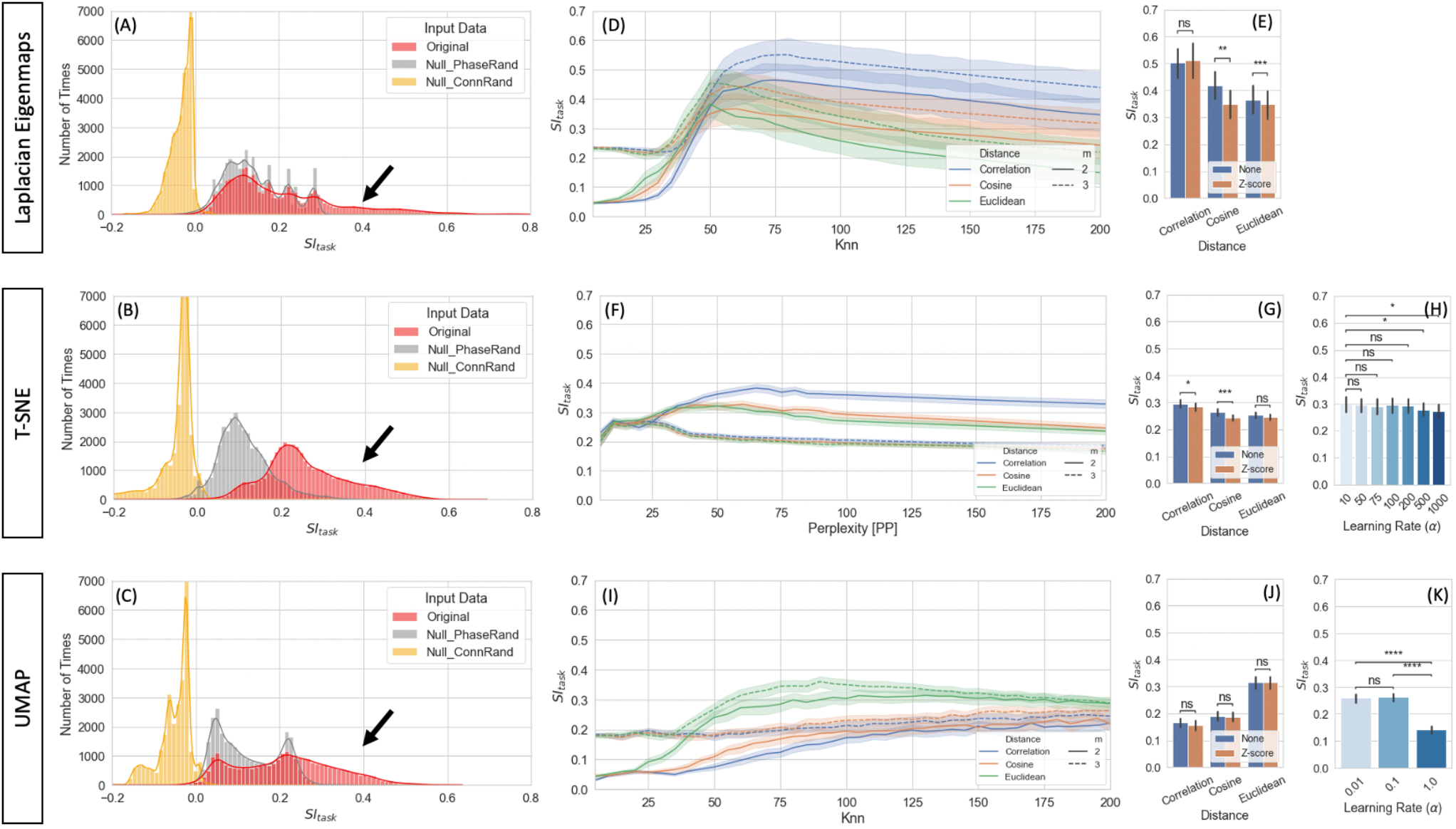
Task separability for single-scan level embeddings. (A) Distribution of *SI*_*task*_ values for original data and null models across all hyperparameters for *LE* for 2D and 3D embeddings *[Total Number of Cases: 20 Subjects * 3 Models * 2 Norm Methods * 3 Distances * 40 Knn values * 2 Dimensions]*. (B) Same as (A) but for *T-SNE*. (C) Same as (A) but for *UMAP*. (D) *SI*_*task*_ for *LE* as a function on distance metric and number of final dimensions for the original data. Bold lines indicate mean values across all scans and hyperparameters. Shaded regions indicate 95% confidence interval. (E) Effects of normalization scheme on *SI*_*task*_ at *K*_*nn*_=75 for original data and the three distances. (F) Same as (D) but for *T-SNE*. (G) Same as (E) but for *T-SNE* and *PP*=75. (H) *SI*_*task*_ dependence on learning rate for *T-SNE* at *PP*=75. (I) Same as (D) but for *UMAP*. (J) Same as (E) but for *UMAP*. (K) Same as (H) but for *UMAP*. In (E,G,H,J & K) Bar height indicate mean value and error bars indicate 95% confidence interval. Statistical annotations: ns = non-significant, * = *p*_*Bonf*_ < 0.05, ** = *p*_*Bonf*_ < 0.01, *** = *p*_*Bonf*_ < 0.001, **** = *p*_*Bonf*_ < 0.0001.

Figure 6.D shows how *SI*_*task*_ changes with distance function and *K*_*nn*_ for *LE* embeddings. Overall, best performance is achieved when using the *Correlation* distance and keeping 3 dimensions. Additionally, *K*_*nn*_ can also have substantial influence on task separability. *SI*_*task*_ is low for *k*_*nn*_ values below 50, then starts to increase until it reaches a peak around *k*_*nn*_*=75* and then decreases again monotonically with *k*_*nn*_. Figure 6.E shows that whether *tvFC* matrices are normalized or not prior to entering *LE* has little effect on task separability.

Figures 6.F-H summarize how task separability varies with distance, perplexity, normalization scheme and learning rate when using *T-SNE*. As it was the case with *LE*, best results were obtained with the *Correlation* distance. We can also observe high dependence of embedding quality with neighborhood size (i.e., perplexity), and almost no dependence with normalization scheme. Regarding learning rate, *SI*_*task*_ monotonically decreases as learning rate increased. Figures 6.I-K show results for equivalent analyses when using *UMAP*. In this case, best performance is achieved with the *Euclidean* distance. Again, we observe high dependence of *SI*_*task*_ with neighborhood size, no dependence on normalization scheme, and a monotonic decrease with increasing learning rate.

Figure 7 shows representative single-scan *2D* embeddings (see Suppl. Fig 4 for *3D* results). First, figure 7.A shows best and worse *LE* embeddings obtained using the *Correlation* distance and *K*_*nn*_ = 25, 75, 125 and 175. Figure 7.B and 7.C shows *2D* embeddings for the same scans obtained using *T-SNE* with the *Correlation* distance and *UMAP* with the *Euclidean* distance respectively. Embedding shape significant differed across *MLTs* and as function of hyperparameters. For *K*_*nn*_=25, all *MLTs* placed next to each other temporally contiguous windows in a “*spaghetti*-like” configuration. No other structure of interest is captured by those embeddings. For *K*_*nn*_ ≥75 (embeddings marked with a green box), those “*spaghetti*” start to break and bend in ways that bring together windows corresponding to the same task independently of whether or not such windows are contiguous in time. If we focus our attention on high quality exemplars (green boxes), we observe clear differences in shape across *MLTs*. For example, *LE* places windows from different tasks in distal corners of the *2D* (and *3D*) space; and the presence of two distinct task blocks is no longer clear. Conversely, *T-SNE* and *UMAP* still preserve a resemblance of the “spaghetti-like” structure previously mentioned, and although windows from both task blocks now appear together, one can still appreciate that there were two blocks of each task. Finally, Figure 7.D & E shows embeddings for the null models at *K*_*nn*_=75 for the best scan. When connections are randomized, embeddings look like random point clouds. When the phase of ROI timeseries is randomized prior to generating *tvFC* matrices, embeddings look similar to those generated with real data at low *K*_*nn*_, meaning they have a “*spaghetti-like*” structure where time contiguous windows appear together, but windows corresponding to the two different blocks of the same task do not.

**Figure 7.**
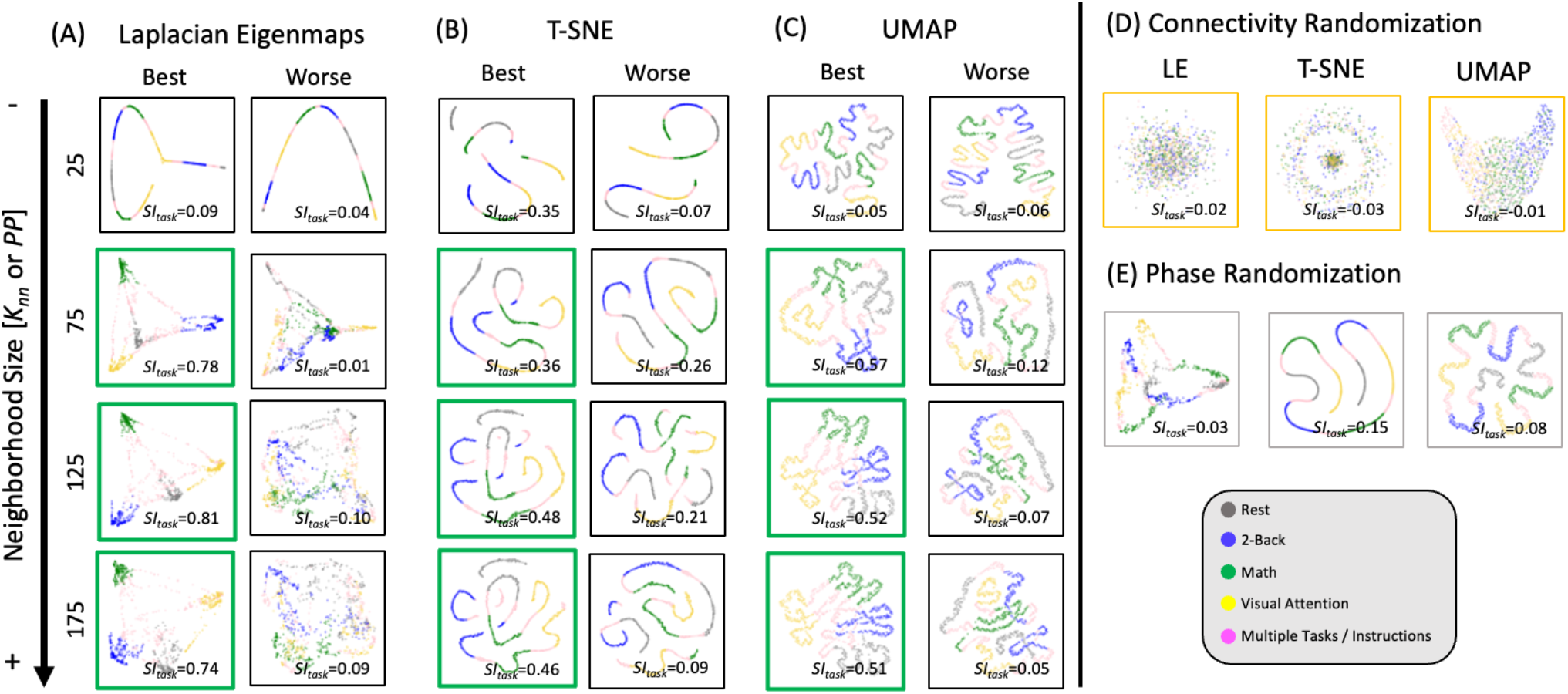
Representative single-scan embeddings. (A) LE embeddings for *K*_*nn*_=25,75,125 and 175 generated with the Correlation distance. Left column shows embeddings for the scan that reached the maximum *SI*_*task*_ value across all possible *K*_*nn*_ values. Right column shows embeddings for the scan that reached worse *SI*_*task*_. In each scatter plot, a dot represents a window of *tvFC* data (i.e., a column in the *tvFC* matrix). Dots are annotated by task being performed during that window. (B) *T-SNE* embeddings computed using the Correlation distance, learning rate=10 and *PP*=25,75,125 and 175. These embeddings correspond to the same scans depicted in (A). (C) *UMAP* embeddings computed using the *Euclidean* distance, learning rate=0.01 and *K*_*nn*_=25,75, 125 and 175. These also correspond to the same scans depicted in (A) and (C). (D) *LE, T-SNE* and *UMAP* embeddings computed using as input the connectivity randomized version of the same scan corresponding to “best” in (A), (B) and (C). (E) *LE, T-SNE* and *UMAP* embeddings computed using as input the phase randomized version of the same scan corresponding to “best” in (A), (B) and (C).

Figure 8 shows clustering evaluation results for group-level embeddings generated with *LE* for the original data. Regarding task separability (Fig. 8.A), the “*Embed + Procrustes*” approach (orange) outperforms the “*Concatenate + Embed*” approach (blue). Importantly, the higher gains for the “*Embed + Procrustes”* approach occur when the transformation is calculated using dimensions beyond three (portion of the orange distribution outlined in dark red in Fig. 8.A). Fig 8.C shows embeddings in which the *Procrustes* transformation was computed with an increasing number of dimensions (from left to right). As the number of dimensions increases towards the data’s *ID*, task separability improves. For example, when all 30 dimensions are used during the *Procrustes* transformation the group embedding show four distinct clusters (one per task), and all subject specific information has been removed (orange histogram in Fig 8.B). Fig. 8.D shows one example of high *SI*_*task*_ for the “Concatenation + Embed” approach. This occurs on a few instances (long right tail of the blue distribution in Fig. 8.B) that corresponds to scenarios where an excessively low *Knn* results in a disconnected graph.

**Figure 8.**
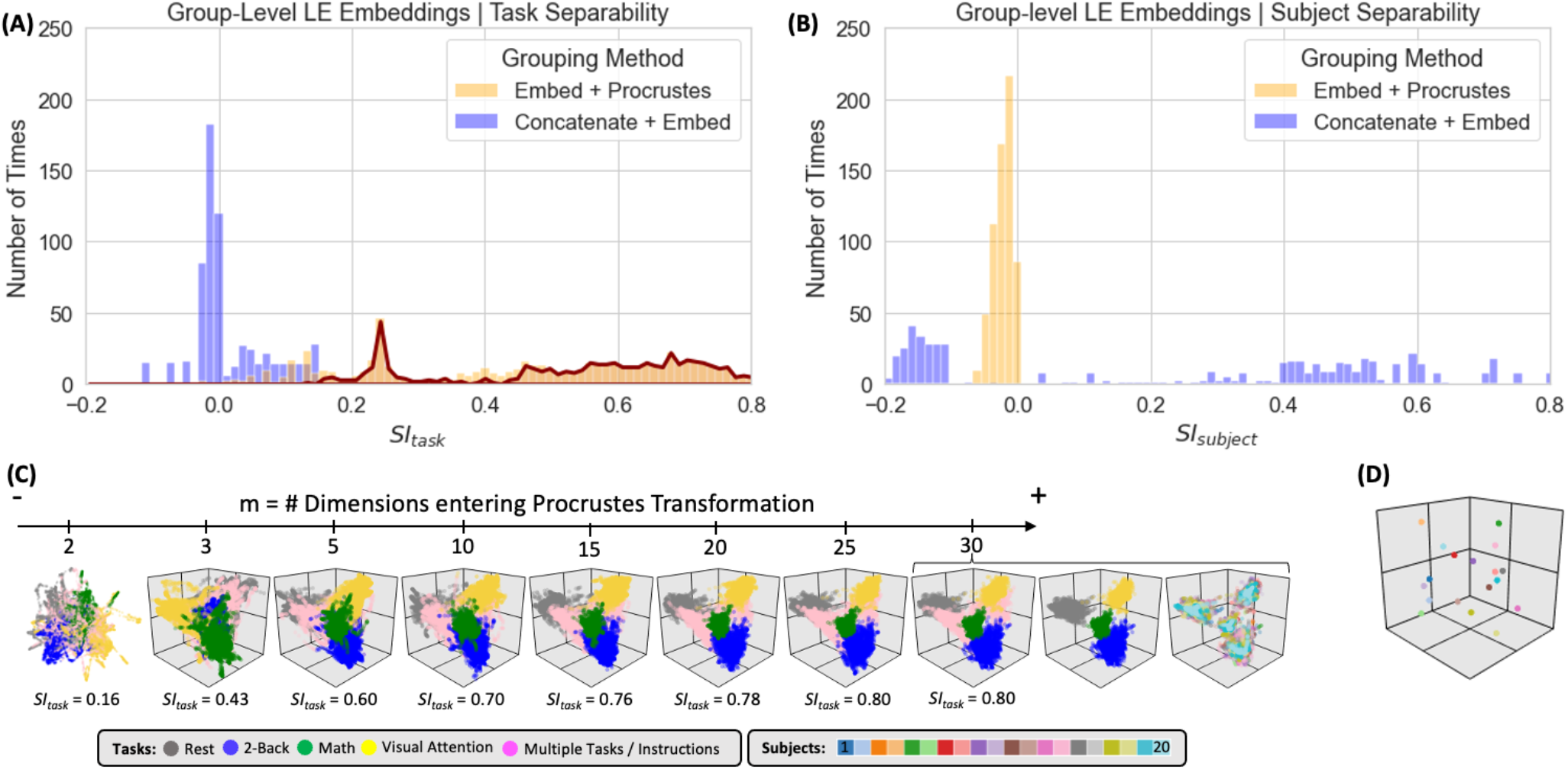
Summary of clustering evaluation for *LE* group embeddings. (A) Histograms of *SI*_*task*_ values across all hyperparameters when using the *Correlation* distance with real data. Distributions segregated by grouping method: *“Embed + Procrustes”* in orange and *“Concatenation + Embed”* in blue. Dark red outline highlights the section of the distribution for “*Embed + Procrustes*” that corresponds to instances where more than 3 dimensions were used to compute the Procrustes transformation. (B) Same as (A) but this time we report *SI*_*subject*_ instead of *SI*_*task*_. (C) Group level *LE* embeddings obtained via “*Embed + Procrustes*” with an increasing number of dimensions entering the Procrustes transformation step. In all instances we show embeddings annotated by task, including windows that span more than one task and/or instruction periods. For m=30 we show two additional versions of the embedding, one in which task inhomogeneous windows have been removed, so that task separability becomes clear, and one when windows are annotated by subject identity to demonstrate how inter-individual differences are not captured in this instance. (D) Representative group embedding with high *SI*_*subject*_ obtained via “*Concatenation + Embed*” approach.

Figure 9 shows clustering evaluation results for *UMAP*. Figure 9.A shows the distribution of *SI*_*task*_ values across all hyperparameters when working with the real data and the *Euclidean* distance. High *SI*_*task*_ values were only obtained when combining scans via the “*Embed + Procrustes*” approach and using more than 3 dimensions during the *Procrustes* transformation (highlighted portion of the orange distribution in Fig 9.A). Fig 9.C shows one example of an embedding with high *SI*_*task*_ computed this way. Clear task separability is observed when annotating the embedding by task (Fig 9.C left). If we annotate by subject identity (Fig 9.C right), we can observe how individual differences have been removed by this procedure. Figure 9.B shows the distribution of *SI*_*subject*_ values. High *SI*_*subject*_ values were only obtained when using the “*Concatenation + Embed*” approach in data that has not been normalized (dark blue outline). Z-scoring scan-level *tvFC* matrices prior to concatenation removes *UMAP* ability to capture subject identity (light blue highlight). Figure 9.D shows a *UMAP* embedding with high *SI*_*subject*_ annotated by task (left) and subject identity (right). The embedding shows meaningful structure at two different levels. First, windowed *tvFC* are clearly group by subject. In addition, for the majority of subjects, the embedding also captures the task structure of the experiment. Results for T-*SNE* group-level embeddings are shown in Suppl. Figure 5. T-*SNE* was also able to generate embeddings that simultaneously capture task and subject information using the “*Concatenation + Embed”* and no normalization. Similarly, it could generate group embeddings that bypass subject identity and only capture subject structure by using the “*Embed + Procrustes*” approach, yet their quality was inferior to those obtained with *UMAP*.

**Figure 9.**
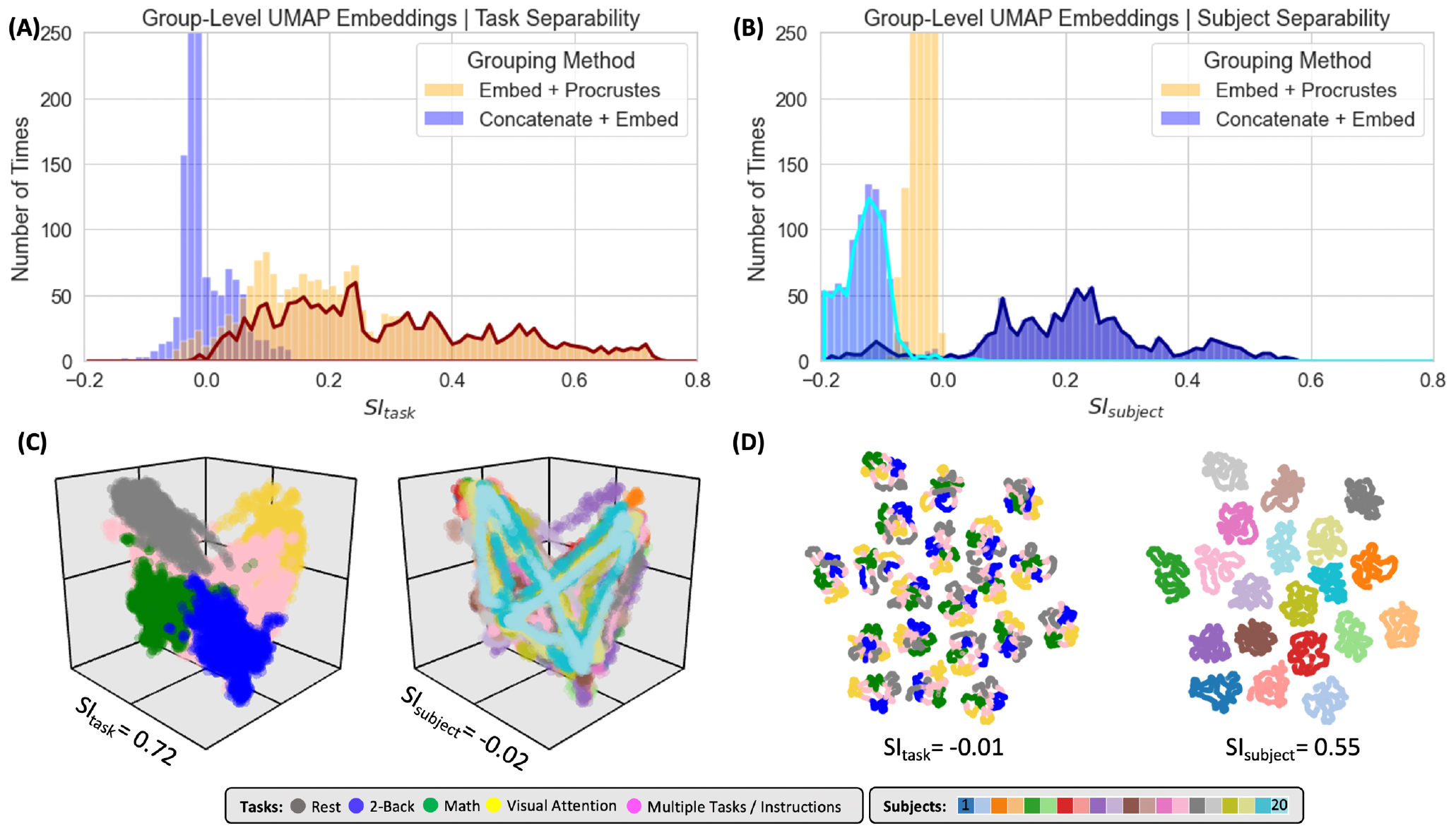
Summary of clustering evaluation for *UMAP* group embeddings. (A) Histograms of *SI*_*task*_ values across all hyperparameters when using *Euclidean* distance on real data. Distributions are segregated by grouping method: “*Embed + Procrustes*” in orange and “*Concatenation + Embed*” in blue. Dark orange outline highlights the section of the “*Embed + Procrustes*” distribution that corresponds to instances where more than 3 dimensions were used to compute the Procrustes transformation. (B) Histograms of *SI*_*subject*_ values across all hyperparameters when using *Euclidean* distance on real data. Distributions are segregated by grouping method in the same way as in (A). Light blue contour highlights the part of the distribution for “*Concatenation + Embed*” computed on data that has been normalized (e.g., Z-scored), while the dark blue contour highlights the portion corresponding to data that has not been normalized. (C) High quality group-level *“Embed + Procrustes”* embedding annotated by task (left) and subject identity (right). (D) High quality group-level “*Concatenation + Embed*” annotated by task (left) and subject identity (right).

### Evaluation for Predictive/Classification Purposes

Figure 10 shows results for the predictive framework evaluation. This evaluation was performed using only embeddings that performed well on the task separability evaluation. For *UMAP*, this includes embeddings computed using the *Euclidean* distance, learning rate = 0.01 and *K*_*nn*_ > 50. For *LE*, this includes embeddings computed using the *Correlation* distance and *K*_*nn*_ > 50. In both instances, we used as input group level embeddings computed using the “*Embed + Procrustes*” aggregation method. We did not perform this evaluation on T-*SNE* embeddings because computational demands increase significantly with the number of dimensions.

**Figure 10.**
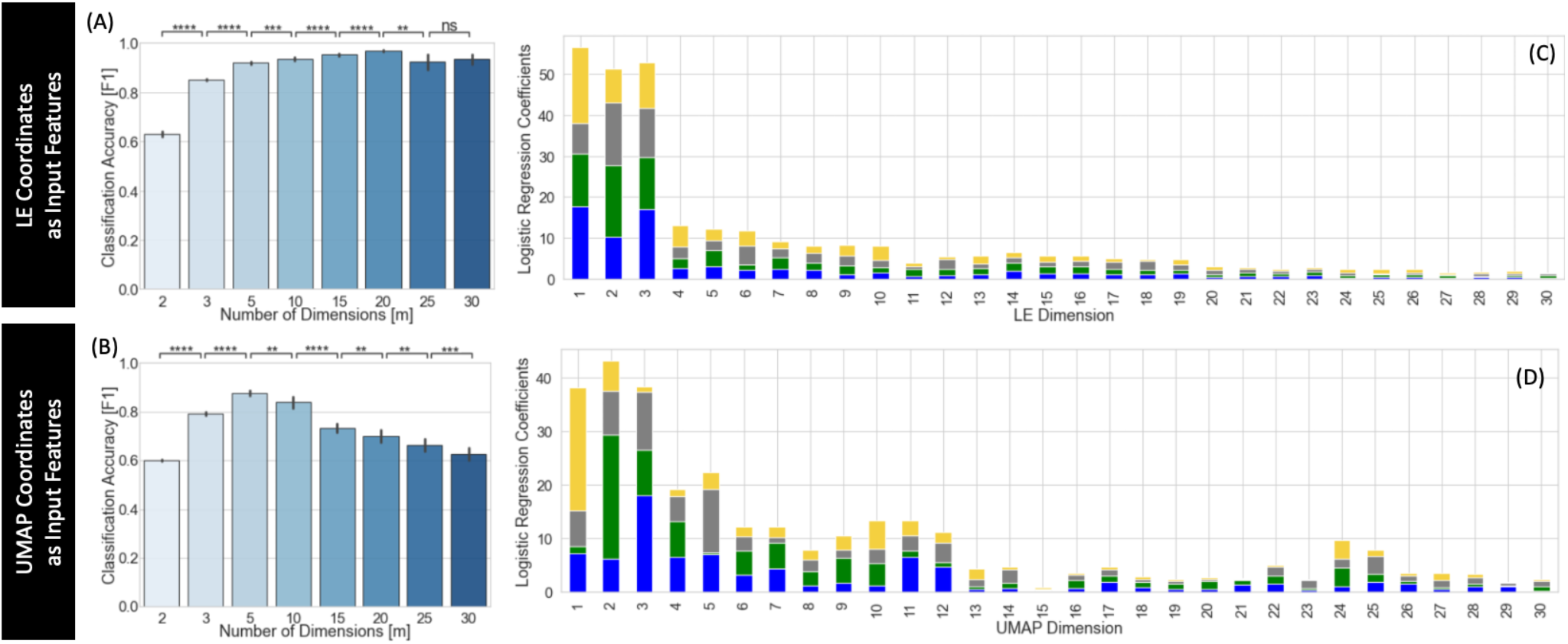
Summary of predictive framework evaluation for LE (A,C) and *UMAP* (B,D) group embeddings. (A) Classification accuracy as a function of the number of LE dimensions used as input features to the logistic regression classifier. Classification improves as m increases up to m = 20. (B) Classification accuracy as a function of the number of UMAP dimensions used as input features to the logistic regression classifier. Classification improves as m increases up to m = 5. Statistical annotations for (A) and (B) as follows: ns = non-significant, * = *p*_*Bonf*_ < 0.05, ** = *p*_*Bonf*_ < 0.01, *** = *p*_*Bonf*_ < 0.001, **** = *p*_*Bonf*_ < 0.0001. (C) Average coefficients associated with each *LE* dimension for classifiers trained using m = 30 dimensions. For each m value, we stack the average coefficients associated with each label, which are colored as follows: blue = 2-back, green = math, yellow = visual attention, grey = rest. (D) Same as (C) but for UMAP.

Figure 10.A shows average classification accuracy for *LE*. Classification accuracy increased significantly with the number of dimensions up to m = 20. Beyond that point, accuracy slightly decreased but remained above that obtained with m ≤ 3. Figure 10.B shows equivalent results for *UMAP*. In this case, accuracy significantly increased up to m=5, but monotonically decreased beyond that point. For m ≥ 15, accuracy was less than that of m=3. Figures 10.C and D shows the average classifier coefficient values associated with each dimension for classifiers trained with m = 30 for LE and UMAP respectively. In both instances we can observe that although the higher contributing features are those for the first three dimensions, there are still substantial contributions from higher dimensions.

## Discussion

The purpose of this work is to understand why, how, and when manifold learning techniques can be used to generate informative low dimensional representations of *tvFC* data. In the theory section, we first provide our argumentation for why we believe it is reasonable to expect *tvFC* data to lie on a low dimensional manifold embedded within the higher dimensional ambient space of all possible pair-wise ROI connections. We next discuss, at a programmatic level, the inner workings of three state-of-the-art manifold learning techniques often used to summarize biological data. This theoretical section is accompanied by several empirical observations: 1) the dimensionality of *tvFC* data is several orders of magnitude lower than that of its ambient space, 2) the quality of low dimensional representations varies greatly across methods and also within method as a function of hyper-parameter selection, 3) temporal autocorrelation, which is prominent in *tvFC* data but not on other data modalities commonly used to benchmark manifold learning methods, dominates embedding structure and must be taken into consideration when interpreting results, 4) while three dimensions suffice to capture first order connectivity-to-behavior relationships (as measured by task separability), keeping additional dimensions up to the *ID* of the data can substantially improve the outcome of subsequent transformation and classification operations.

### Intrinsic Dimension

FC matrices are often computed using brain parcellations that contain somewhere between 100 and 1,000 different regions of interest. As such, *FC* matrices have dimensionality ranging from 4,950 to 499,500.

Here we used a parcellation with 157 regions, resulting in 12,246 dimensions. Our results indicate that *tvFC* data only occupies a small portion of that immense original space, namely that of a manifold with dimensionality ranging somewhere between 3 to 25. This suggests that *tvFC* can be aggressively compressed with little loss in information. That said, such low *ID* is not evidence of slow dynamics or implies that the brain only took a handful of *FC* configurations. For example, let’s consider a latent space where each dimension ranges between -1 and 1 (as it is the case with Pearson’s correlation) in increments of 0.2 (e.g., -1.0, -0.8, -0.6 … 0.6, 0.8. 1.0). As such, there are only 11 possible values per dimension. A hypothetical 3D space with dimensions defined that way could represent up to 11^3^ = 1331 different *FC* configurations. If we consider 5 dimensions, the number of configurations goes up to 1.6e^5^. With 25 dimensions, we reach 1.1e^26^ different possible FC configurations to be represented in such space. In other words, low *ID* should not be regarded as a sign of *FC* being relatively stable, but as confirmatory evidence that *FC* patterns are highly structured and in agreement with prior observations such as the fact that the brain’s functional connectome adhere to a very specific network topography (e.g., small world network (Sporns and Honey, 2006)) or that task performance does not lead to radical *FC* re-configurations (Cole et al., 2014; Krienen et al., 2014).

Using the *twoNN* estimator, we found that the *ID*_*global*_ of *tvFC* from rest periods is significantly higher than that of the other three tasks (Fig 5.C), suggesting that *FC* traverses a larger space of possible configurations during rest compared to when subjects are engaged in tasks with more precise cognitive demands. This agrees with prior work suggesting an overall increase in the stability of *FC* for task as compared to rest (Billings et al., 2017; Chen et al., 2015; Elton and Gao, 2015; Liu and Duyn, 2013). It also suggests that *ID* can be a valuable summary statistic for *tvFC* data. In machine learning, *ID* is used to evaluate the complexity of object representations at different levels of deep neural networks (Ansuini et al., 2019) and their robustness against adversarial attacks (Amsaleg et al., 2017). In biomedical research, *ID* has also been used to characterize the amount of variability present in protein sequence evolution (Facco et al., 2019), and to explain why humans can learn certain concepts from just a few sensory experiences (i.e., “few-shot” concept learning; (Sorscher et al., 2022)). Given that several psychiatric conditions and neurological disorders have been previously associated with altered functional connectivity dynamics (Damaraju et al., 2014; Fiorenzato et al., 2018; Kaiser et al., 2019; Rashid et al., 2016), future research should evaluate the value of *ID* as a marker of clinically relevant aberrant *tvFC*.

Finally, *ID* for *tvFC* data was estimated to be near to the number of dimensions that can be easily visualized (i.e., 2 or 3 dimensions). This explains the success of *MLTs* at summarizing *tvFC* data reported here and elsewhere (Gao et al., 2021; Gonzalez-Castillo et al., 2019; Rué-Queralt et al., 2021). Yet for two *ID* estimators (*lPCA* and *twoNN*), *ID* was estimated to be greater than 3 (Figure 5.A-B). This suggests that one ought to keep and explore additional dimensions up to *ID* whenever possible. We empirically demonstrate the value of keeping additional dimensions up to *ID* in two scenarios: across-scan embedding alignment and task classification. For example, in figure 8.C we show how task separability for *Procrustes*-based group embeddings substantially improves when more than three dimensions are used to compute the transformation. The *Procrustes* transformation has several applications in fMRI data analysis including hyper-alignment of voxel-wise responses (Haxby et al., 2011), alignment of macro-anatomical functional connectivity gradients (Margulies et al., 2016) and generation of bidirectional mappings between fMRI responses and natural language description of scenes in naturalistic experiments (Vodrahalli et al., 2018). The benefits of using dimensions beyond those being interpreted during alignment has been previously reported for functional gradients by McKeown and colleagues (2020) who showed that alignment towards a template space significantly improves when using 10 dimensions instead of three (the number of gradients often explored and interpreted in studies that rely on this technique (Hardikar et al., 2022; Margulies et al., 2016; Mckeown et al., 2020; Tian et al., 2020)). The supplementary materials from that study (Suppl. Figure 2 in (Mckeown et al., 2020)) show that by keeping the additional seven dimensions the authors approximately doubled the amount of variance explained available as input to the *Procrustes* transformation, yet no clear heuristic was provided about how to select the optimal number of inputs. Our results suggest that *ID* could help generate such heuristics and help inform how many dimensions ought to be explored and retained during analysis.

### Hyper-parameter selection

Our results demonstrate that although *MLTs* can create low dimensional representations of *tvFC* data that capture structures of interest at different levels (i.e., subject identity, mental state), hyper-parameter selection was critical to their success. Particularly important was the selection of distance function and neighborhood size for single-scan embeddings. For group embeddings, aggregation method and normalization scheme also played a critical role.

One common theme across explored *MLTs* is the construction of a neighborhood graph in early stages of the algorithm (e.g., Fig 2.D). The final form of such graph depends, to a large extent, on how one decides to quantitatively measure dissimilarity between connectivity patterns (distance function) and how big one expects neighborhoods of interest to be (*K*_*nn*_ or *PP*). For *LE* and *T-SNE* best results were obtained using *Correlation* distance, which measures the degree of linear relationship between two sets of observations. *Correlation* is often used to quantify similarity in fMRI data, whether it be between timeseries (Biswal et al., 1995; Finn et al., 2014), activity maps (Matsui et al., 2022; Yarkoni et al., 2011) or connectivity matrices (Cole et al., 2014). Therefore, *Correlation’s* ability to meaningfully quantify similarity is well accepted and validated in the field. Moreover, previous work with non-imaging data suggest that the *Euclidean* distance fails to accurately capture neighborhood relationships in high dimensional spaces (Beyer et al., 1999) due to the curse of dimensionality, and that the *Correlation* distance is more appropriate for clustering and classification problems on high dimensional data (France and Carroll, 2009). Our results confirm that is also the case for *tvFC* data. One exception was *UMAP*, which performed best with the *Euclidean* distance for mid-size neighborhoods and became successively more equivalent to the other distances as neighborhood size increased. As described on the theory section, *UMAP* contains a distance normalization step (Fig 4.E) aimed at ensuring that each sample is connected to at least one other sample in the neighboring graph. We believe this step is the reason why the *Euclidean* distance outperforms the *Correlation* distance in *UMAP*. First, as discussed in the original description of the algorithm by McInnes et al. (2018), this normalization step increases robustness against the curse of dimensionality and, as such, it helps mitigate some of the undesired outcomes of using *Euclidean* distances with high dimensional data. Second, because *Correlation* distances are in the range [0,2], *rho* (Eq. 12) always takes values near zero (Supplementary Figure 7.A) and *sigma* (Eq. 13) is restricted to a narrow range of values (Supplementary Figure 7.C). These two circumstances contribute to worsening UMAP’s ability to capture global structure (e.g., same task in temporally distant blocks) when using the *Correlation* distance.

In summary, when using *MLTs* and *tvFC* data one ought to attempt to mitigate the negative effects derived from the curse of dimensionality—namely the fact that high dimensional spaces tend to be sparse and *Euclidean* distances become progressively meaningless—either by the selection of an alternative distance metric (e.g. *Correlation*) or by relying on algorithms with some built-in level of protection against it (e.g., *UMAP*).

Neighborhood size, the second key parameter for graph generation, can be thought of as a way of setting the scale at which information of interest is expected. In our test data, subjects engaged with four different tasks during two temporally disconnected 180s blocks. Given our sliding window parameters (*Window Duration* = 45s; *Window Step* = 1.5s) this results in 91 windows per task block and a total of 182 windows per task on each scan. For all methods, we observed bad task separability (low *SI*_*task*_) at the smallest neighborhood sizes (e.g., < 60). This is because setting such small values often precluded the graph to capture neighboring relationships beyond those due to large overlap in the number of samples contributing to temporally contiguous windows. As we approach neighborhood values above 70, we start to observe the best *SI*_*task*_ values. This is because at this point, the graph can now capture neighborhood relationships between windows corresponding to different blocks of the same task and embeddings start to show structure and clusters that relate to the task. As neighborhood size keeps increasing, *SI*_*task*_ slowly degrades because more windows from different tasks end up being marked as neighbors during the construction of the graph.

### Challenges for resting-state fMRI

In the previous paragraph, we were able to explain the relationship between task separability and neighborhood size because we know the scale of the phenomena of interest and have labels for tasks. But what about situations where such information is missing? For example, should we decide to use *MLTs* to explore the dynamics of *tvFC* during resting state, what is the recommended neighborhood size to use? This is quite a difficult question. Initially, one could opt to rely on existing heuristics from machine learning such using a neighborhood size equal to the square root of the number of samples (Hassanat et al., 2014), but such heuristic would have resulted in a value of 27, which is far from optimal. Similarly, using default values in existing implementations would have also proven sub-optimal here (e.g., *PP* = 30 in *scikit*-*learn* implementation of T-*SNE*). A second approach would be to fine tune neighborhood size using some hyper-parameter optimization scheme, yet those methods require larger datasets and an objective function to optimize. These two requirements are hard to meet when working with *tvFC* data. First, in contrast with other data modalities such as natural images, speech or genomics, fMRI datasets are often of a limited size (although this is changing thanks to recent international efforts such UK’s Biobank (Miller et al., 2016)). Second, defining an objective function in this context is quite challenging. Not only it requires labeled data—which is almost inexistent for resting-state—but as our data shows, it can be misleading (Fig. 9.D shows an embedding that captures both subject and task identity but has low *SI*_*task*_). A third approach would be to transfer heuristics from studies such as this one. We recently took this approach in a study looking at the temporal dynamics of functional connectivity during rest. We used the same multi-task dataset evaluated here to inform our selection of *K*_*nn*_ for *LE*. Using this approach, combined with reverse inference via *Neurosynth* (Yarkoni et al., 2011), we were able to show that resting-state *tvFC* patterns sitting at the corners of *LE* embeddings correspond to mental activities commonly reported as being performed during rest (Gonzalez-Castillo et al., 2019). Finally, an additional alternative would be to use newer versions of the algorithms that do not require a priori selection of neighborhood size such as perplexity-free *T-SNE* (Crecchi et al., 2020) or optimize concurrently at several scales (Lee et al., 2015), yet the performance of those algorithmic variants in *tvFC* data should be explored first.

### Group-level Aggregation

Functional connectivity is characterized by large inter-subject variability; and subject identification across sessions is possible using both static (Finn et al., 2014) and time-varying (Betzel et al., 2022) *FC* patterns as a form of fingerprinting. Here, inter-subject variability is clearly captured by group embeddings computed using the “*Concatenate + Embed*” approach on non-normalized data (see Figures 8.D and 9.D). If data is normalized prior to concatenation, then subject identity is no longer depicted in the embeddings (see Figures 8.C and 9.C). This suggests that one key differentiating aspect across subjects is differences in the mean and/or standard deviation of ROI-to-ROI functional connectivity traces. Supplementary Figure 6.A-B shows how the distributions of these two summary metrics vary across subjects, and how those differences are removed by the normalization step (Suppl. Figure 6.C-D).

The second way to remove subject identifiability from the group-level embeddings is to generate those using the “*Embed + Procrustes*” approach. This works well independently of whether data is normalized or not. The fact that the *Procrustes* transformation—which only includes translations, rotations and uniform scaling—can bring scan-level embeddings into a common space where windows from the different tasks end up in the same portion of the lower dimensional space suggest that scan-level embeddings share a common geometrical shape and therefore that within-subject relationships between the *FC* patterns associated with the four different tasks are largely equivalent across subjects.

### The role of Temporal Autocorrelation

While the “*connectivity randomization*” null model always resulted in embeddings with no discernable structure (Figure 7.D), that was not the case for the “*phase randomization*” null model (Figure 7.E). Although both models remove all neuronally meaningful information, they differ in one critical way. In the *“connectivity randomization”* model, randomization happens after the construction of the *tvFC* matrix. Conversely, in the *“phase randomization”* model, randomization is applied over the ROI timeseries and therefore precedes the sliding window procedure. Because of this, a substantial amount of temporal autocorrelation is reintroduced in this second null model during the sliding window procedure. This results in *FC* patterns from temporally contiguous windows being very similar to each other, even if those patterns are neuronally and behaviorally meaningless. When *MLTs* are then applied to this surrogate data, those temporally contiguous windows appear in proximity. Moreover, because each individual task block spans 91 such windows, one could get the impression that embeddings computed on this second null model were able to recover some task structure. Yet, that is not the case. All they do is to recapitulate the time dimension. They never place together windows from separate task blocks as it is the case with the embeddings computed over real data. In summary, the results from the “*phase randomization*” model do not suggest *MLTs* will show good task separability in data with no neuronally driven *FC*. What they highlight is the importance of considering the role of temporal autocorrelation when interpreting and evaluating embeddings.

Because the goal of *MLTs* is to preserve local structure over global structure, and temporal autocorrelation is the largest source of local structure in *tvFC* data obtained with sliding window procedures, embeddings will easily recapitulate the time dimension in such data. Also, as mentioned in the discussion about the role of normalization in group-level embeddings above, *MLTs* tend to separate *FC* snapshots that have different mean and/or standard deviation. These two observations should be considered when selecting *FC* datasets for *MLT* benchmarking or interpreting embeddings generated with them. For example, benchmarking datasets should always include multiple temporally distant repetitions of each phenomenon of interest (e.g., mental states). The minimum temporal separation between them should be larger than the intrinsic temporal autocorrelation properties of the data. Moreover, to minimize systematic shifts in mean and standard deviation, such repetitions should occur within the confines of individual scans. In this sense, we believe that long multi-task scans acquired as subjects perform and transition between different tasks or mental states that repeat on several distant occasions might be the optimal type of data for benchmarking *MLTs* and related methods on *tvFC* data. Similarly, when using *MLTs* for summarization and interpretation of *tvFC*, one ought to ensure that observations are not easily explained by the two confounds discussed here: temporal autocorrelation induced during the generation of *tvFC* traces and/or systematic differences in average or volatility values due to factors such as using data from different subjects or different scans.

### Heuristics and Future work

One goal of this work was to provide a set of initial heuristics for those looking to apply *MLTs* to *tvFC* data. The following recommendations emerge from our observations. First, while all three evaluated *MLTs* can generate meaningful embeddings, they showed different behaviors. Overall, *LE* resulted in the best task separability. This occurred when using the *Correlation* distance and *K*_nn_ greater than 50. Although embedding quality is modulated by *K*_nn,_ and the optimal *K*_*nn*_ will be dataset specific, our results seem to suggest that in general the use of larger values is safer as it helps avoid disconnected graphs and embeddings that only capture inter-scan or inter-subject differences. Two additional benefits of *LE* are low computational demands, no optimization phase (which means no additional hyper-parameters to choose from). Additionally, previous applications of *LE* to *FC* data have proven quite successful (Gonzalez-Castillo et al., 2019; Margulies et al., 2016; Mckeown et al., 2020; Rué-Queralt et al., 2021). As such, *LE* might be a good choice for initial choice for those willing to start using *MLTs* on *tvFC* data.

That said, *LE* was the only method that was not able to simultaneously capture two different levels of information (i.e., task and subject identity). In this regard, *T-SNE* and *UMAP* outperformed *LE*, and therefore if one seeks to obtain such multi-scale representations, these two methods may constitute a better alternative. Between both methods, *UMAP* is initially a better candidate because of its computational efficiency. This is particularly true if one is willing to explore dimensions beyond three, as T-SNE’s computing times becomes significantly larger as the number of required dimensions increase. Finally, for *UMAP* our results suggest the use of the *Euclidean* distance and a preference over larger *K*_*nn*_ values in the same manner as just discussed for *LE*.

Finally, it is worth noting that our exploration of *MLTs* is by no means comprehensive. This work is limited not only in terms of data size (*20* scans) and evaluation metrics (clustering and classification), but also in terms of the breath of methods being evaluated. For example, manifold estimation can also be accomplished via multidimensional scaling (Kruskal, 1964), *ISOMAP* (Tenenbaum et al., 2000), diffusion maps (Coifman et al., 2005) or T-PHATE (Busch et al., 2022), to name a few additional MLTs not considered here. All these other methods have been previously applied to fMRI data using either regional levels of activity (Busch et al., 2022; Gao et al., 2021) or static FC (Ioannis K Gallos et al., 2021; Ioannis K. Gallos et al., 2021) as inputs. Future research should evaluate their efficacy on *tvFC* data. Meaningful dimensionality reduction can also be accomplished in other ways such as linear decomposition methods (e.g., Principal Component Analysis, Independent Component Analysis, Non-negative Matrix Factorization, etc.), using deep neural networks (e.g., autoencoders (Wang et al., 2014)) or TDA methods (e.g., Mapper (Saggar et al., 2018)). All these alternatives should also be considered as valuable candidates for dimensionality reduction of *tvFC* data. Of particular interest for resting-state applications is the case of autoencoders because evaluation of low dimensionality representations in this case do not necessarily require labeled data, and prior work has shown their ability to capture generative factors underlying resting-state activity (B.-H. Kim et al., 2021; J.-H. Kim et al., 2021).

## Conclusions

Dimensionality reduction, particularly manifold learning, can play a key role in the summarization and interpretation of *tvFC* data, especially when such data is utilized to study experimentally unconstrained phenomena such as mind wandering, spontaneous memory recall and naturalistic paradigms. Yet, because most *MLTs* are benchmarked and developed using data modalities with different properties to that of *tvFC*, extreme caution must be exerted when transferring methods and heuristics from these other scientific disciplines. To alleviate this issue, here we evaluated three state-of-the art *MLTs* using labelled *tvFC* data. This evaluation suggests that *LE* and *UMAP* outperform T-*SNE* for this type of data. It also highlights the confounding role of temporal autocorrelation, and how it can artifactually inflate evaluation metrics. While this report is limited, we hope it actively contributes to the steady building of a much-needed bridge between the fields of neuroscience and machine learning. Future steps in this direction should include the generation of neuroimaging-based benchmarking datasets that can be easily added to existing benchmarking efforts (Campadelli et al., 2015), and the development of *MLTs* tailored to address the specific needs and characteristics of *tvFC* data.

## Acknowledgements

This research was also possible thanks to the support of the National Institute of Mental Health Intramural Research Programs (ZIAMH002783, ZICMH002968). Portions of this study used the high-performance computational capabilities of the *Biowulf* Linux cluster at the National Institutes of Health, Bethesda, MD (*biowulf.nih.gov*).

## Footnotes

1 Some implementations of the *LE* algorithm, like the one in *skicit-learn*, work with the normalized version of the *Laplancian* matrix.

## Notes

### Competing Interest Statement

The authors have declared no competing interest.

