## Supplementary Materials for "Manifold Learning for fMRI time-varying FC"

### Supplementary Note 1 – Intrinsic Dimension

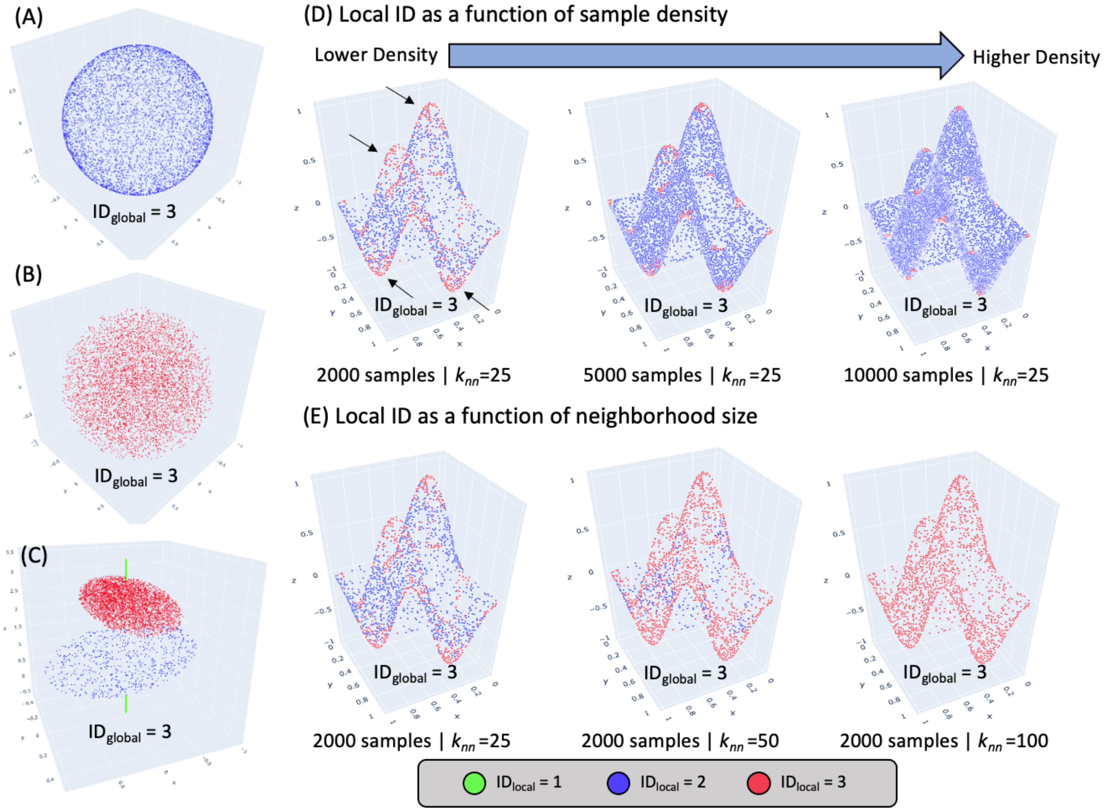


Supplementary Figure 1. (A) Dataset with all samples lying on the surface of a 3D sphere. (B) Dataset with all samples spanning the volume delimited by a 3D sphere. (C) Dataset with varying levels of IDlocal as samples sit on a line, inside a 2D ellipsis, or span the volume of a 3D ellipse. (D) Dependence of $ID_{local}$ on sampling density. $ID_{local}$ is estimated on three different versions of the same dataset with varying levels of sampling density and using a fixed neighborhood size of $k_{nn}$=25 samples. (E) Dependence of $ID_{local}$ on neighborhood size used during its estimation. All panels show the same dataset, yet $ID_{local}$ estimates were obtained using three different neighborhood sizes ranging from 25 to 100 samples. In all plots, data samples are colored according to their $ID_{local}$ estimate values.

Supplementary figure 2.A-C shows three toy datasets with the same $ID_{global}$ but different $ID_{local}$. Pnael A shows a dataset where all samples lie on the surface of a 3D sphere. The $ID_{global}$ for this dataset is 3, and $ID_{local}$ equals 2 for all samples. Panel B shows a second dataset with $ID_{global}$ equal to 3; yet this time, samples are evenly distributed over the volume inside the 3D sphere, and their $ID_{local}$ is 3. Finally, Panel C shows a dataset with varying $ID_{local}$ such that one portion of the data lies on a line ($ID_{local}$ =1; green samples), a second portion sits on a 2D ellipsis ($ID_{local}$ =2; blue samples), and the third portion spans the volume delimited by a 3D ellipsis ($ID_{local}$=3; red samples). Despite those differences in $ID_{local}$, the $ID_{global}$ for this third dataset is also 3. Next, panels D through E illustrate $ID_{local}$ estimates dependence on data density and $k_{nn}$ on a fourth dataset composed of points lying on a 2D plane with four protruding peaks (black arrows). The $ID_{global}$ for this dataset is also 3. The $ID_{local}$ is 2 for most samples except those located at the inflection points where the protuberances emerge from the horizontal plane or reach their respective maximum and minimum values. Yet, $ID_{local}$ estimates not always convey such structure. For example, the lower the density, the larger the number of samples identified as having $ID_{local}$=3 (Panel D). This is because as density decreases, the diameter for a vicinity of fixed size (e.g., $k_{nn}$=25) increases, and with it the likelihood of including samples near or at inflection points. Similarly, given a fixed density, as $k_{nn}$ increases so it does the number of samples with $ID_{local}$=3. This is also due to the increase in diameter. In summary, *ID* estimates are not solely dependent on the structure of the data itself, but also on other parameters such as sampling density, neighborhood size and noise. These additional factors must be considered when interpreting results.

#### Supplementary Note 2 – Distance Functions

For a distance function $d(x, y)$ to be a metric in $\mathbb{R}^N$, $d(x, y)$ must have three properties:

1. Identity: $d(x, y) = 0 \; iff \; x = y$
2. Symmetry: $d(x, y) = d(y, x)$
3. Triangular inequality: $d(x, y) \leq d(x, z) + d(z, y)$

Although desirable, the manifold learning algorithms discussed in this work can work with distance functions that do not fulfill all properties of a metric.

##### Euclidean Distance

The Euclidean distance between two points (or random variables) $x = \{x_1, x_2, .. x_N\}$ and $y = \{y_o, y_1, .. y_N\}$ in a *N* dimensional space $\mathbb{R}^N$ is the length of the line segment going between them. It is given by:

$$d_{Eucl}(x, y) = \sum_{d=1}^{N} (x_d - y_d)^2 \text{ (Eq. 1)}$$

The Euclidean distance is zero for points sitting on the same location. It is greater than zero otherwise. There is no maximum limit for how large the Euclidean distance can be. The Euclidean distance satisfy all properties of a proper distance metric. That said, it is worth noticing that the Euclidean distance tends to increase monotonically with the dimensionality of the space. For this reason, the Euclidean distance is not always recommended when dealing with high dimensional data such as *tvFC* data.

##### Correlation Distance

The correlation distance between two random variables $x = \{x_1, x_2, .. x_N\}$ and $y = \{y_o, y_1, .. y_N\}$ that belong to an N dimensional space $\mathbb{R}^N$ is given by:

$$d_{Corr}(x, y) = 1 - r(x, y) \text{ (Eq. 2)}$$

where $r(x, y)$ is the Pearson's correlation between $x$ and $y$. $d_{Corr}(x, y)$ ranges between zero and two. Because it is based on the Pearson's correlation, $d_{Corr}(x, y)$ measures dissimilarity in terms of the strength of a putative linear relationship between the variables. Importantly, because $r(x, y)$=1 if and only if there is $a > 0$ and $b \in \mathbb{R}$, it is possible for $d_{Corr}(x, y)$ to be zero despite $x$ and $y$ not being equal. In other words, $d_{Corr}(x, y)$ is shift- and scale-invariant, and consequently it does not initially fulfill the identity property of metrics. One way to ensure the fulfillment of this property is to restrict its application to the space of normalized random variables (i.e., those with zero mean and unit variance).

Regarding symmetry, $d_{Corr}(x, y)$ is symmetric because $r(x, y)$ is also symmetric. Finally, $d_{Corr}(x, y)$ does not fulfil the triangular inequality property, and it cannot be considered a metric.

#### Cosine Distance

The cosine distance measures distance between points in terms of the angle (θ) between them. It is given by the following formula:

$$d_{Cos}(x, y) = 1 - cos(\theta) \text{ (Eq. 3)}$$

The minimum cosine distance is zero and occurs when θ equals zero degrees. The maximum cosine distance is two and occurs when θ is 180 degrees. A cosine distance of 1 happens both for θ equal 90 and 270 degrees. As it is the case with the Correlation distance, the Cosine distance is also not a "proper" metric because it cannot fulfill the identity and triangular inequality requirements. Additionally, one must be aware that the angle between two points depends on where the origin sits. Pre-processing steps that shift the location of the origin (e.g., mean removal) will affect estimates of dissimilarity based on the cosine distance. That is not the case for the other two distance functions.

### Supplementary Note 3 – Perplexity

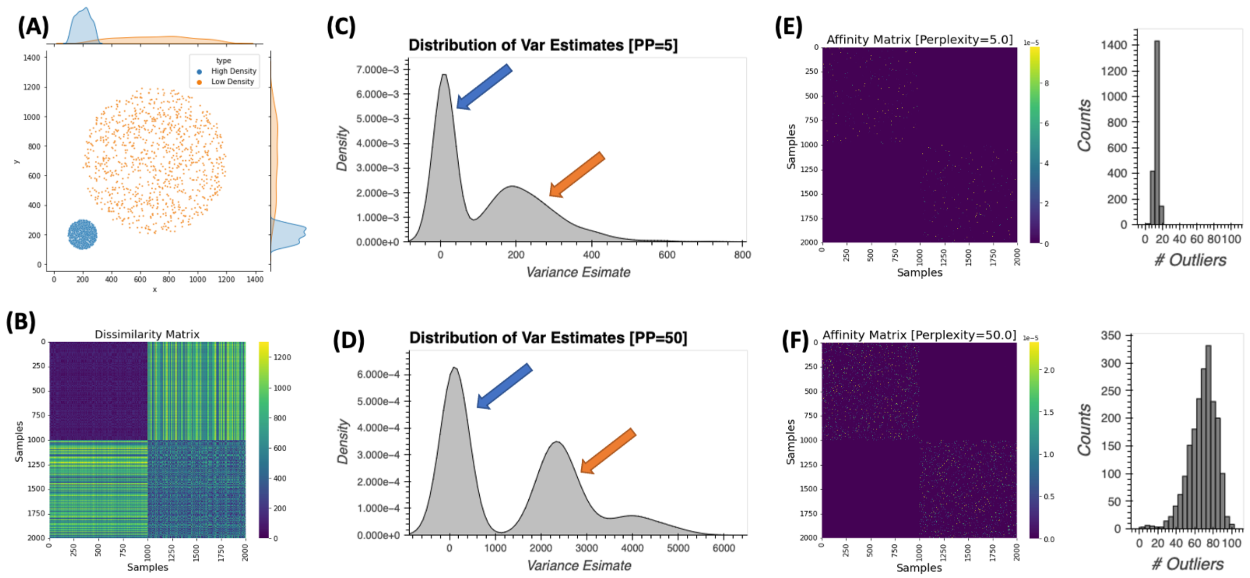


Supplementary Figure 2. Relationship between perplexity and neighborhood size in *T-SNE*. (A) Simulated dataset with two different densities (light blue = high density, orange = low density). (B) Dissimilarity matrix for the simulated dataset (d= Euclidean distance). (C) Distribution of sample-wise estimates of variance for Gaussian kernels centered at each sample for a perplexity of 5. (D) Same as (C) but for a perplexity of 50. (E) *T-SNE* affinity matrix for perplexity of 5. Next to the affinity matrix there is a histogram with counts of neighbors per sample (with neighbors defined as those with substantially larger affinity). (F) Same as (E) for perplexity = 50. Code to generate this figure in Notebook (*S02_Perplexity_SuppFig02*)

The perplexity (*PP*) of a probability distribution (*p*) is a measure of how well *p* summarizes the data. Mathematically it is defined as:

$$PP = 2^{H(p)} \quad \text{(Eq. 4)}$$

where $H(p)$ is the entropy of the distribution. In *T-SNE*, the perplexity lets the user define the scale at which they want to explore the data. Internally, it is used to compute estimates of the variance of the Gaussian distributions that are used to generate the affinity matrix of the data in original space. By setting a common *PP* across all distributions centered at different points, *T-SNE* ensures those distributions are equivalent in terms of the number of samples they embrace, even if density of the input data is not constant across the sample.

Supplementary Figure 2 exemplifies this process. Panel A shows a simulated dataset with 2000 samples. Half the samples are centered around position (x=200, y=200) and have high local density (blue samples are quite closed to each other). The other half of the dataset is centered around position (x=700, y=700) and has a lower local density (orange samples). Panel B shows the dissimilarity matrix for this dataset.

Panel C shows the distribution of variance estimates for Gaussian kernels centered at each sample when working with a perplexity equal to 5. We observe a binomial distribution with peaks centered approx. at $\widehat{\sigma_i^2} = 10$ and $\widehat{\sigma_i^2} = 230$ (colored arrows). The left peak (blue arrow) corresponds to variance estimates for samples in the high-density cloud (blue samples). The right peak (orange arrow) corresponds to variance estimates for samples in the low-density cloud (orange samples). To maintain a constant perplexity across the sample (*PP*=5), higher variance is needed for the kernels set on samples with lower density.

Next, panel D shows the same results for perplexity equal 50. We also observe a binomial distribution, but this time peaks are centered approx. at $\widehat{\sigma_\iota^2} = 110$ (blue arrow) and $\widehat{\sigma_\iota^2} = 2750$ (orange arrow). This second simulation exemplifies how, for the same sample, higher perplexity will translate into higher variance estimates.

Finally, panel E shows T-SNE affinity matrix for data in original space obtained when PP=5. We can observe that each row only contains a few points with high affinity. Next, to the affinity matrix we plot a histogram with those counts. On average, each point has approximately 10 other points with high affinity (i.e., points considered to be close neighbors). Panel F shows equivalent results when PP=50. Now the average number of neighbors have increases to 68. These results show the strong relationship between perplexity and neighborhood size in *T-SNE*.

#### Supplementary Note 4 – UMAP Minimum Distance

Supplementary Figure 3 shows the effect of *UMAP* hyper-parameter minimum distance (*min_dist*) on the final embeddings. We plot embeddings for six different executions of the *UMAP* algorithm on the same data using the same hyper-parameters (*d*=Euclidean, $k_{nn}$=90, *m*=2, $N_{epoch}$=500, $\alpha_{init}$=0.01) with the only difference being *min_dist* which we varied between 0.1 and 0.95 in steps of 0.15. The overall shape of the embedding is the same across all six panels and in all instances the task structure was preserved. As minimum distance increases the separation between the closest samples increases accordingly.

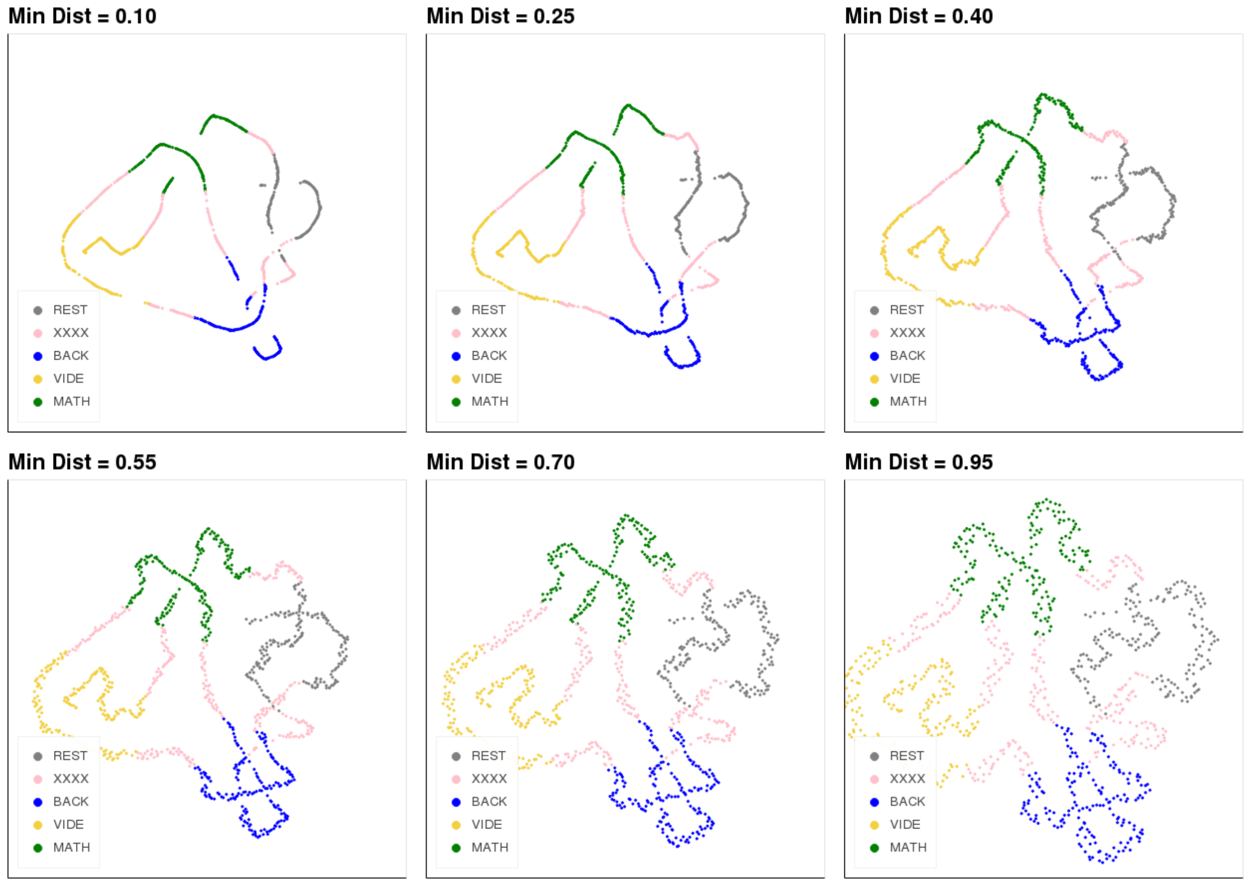


Supplementary Figure 3. UMAP results when the hyper-parameter minimum distance varies between 0.10 and 0.95 in intervals of 0.15. The higher the minimum distance, the farther apart closest points are from each other.

### Additional Supplementary Figures

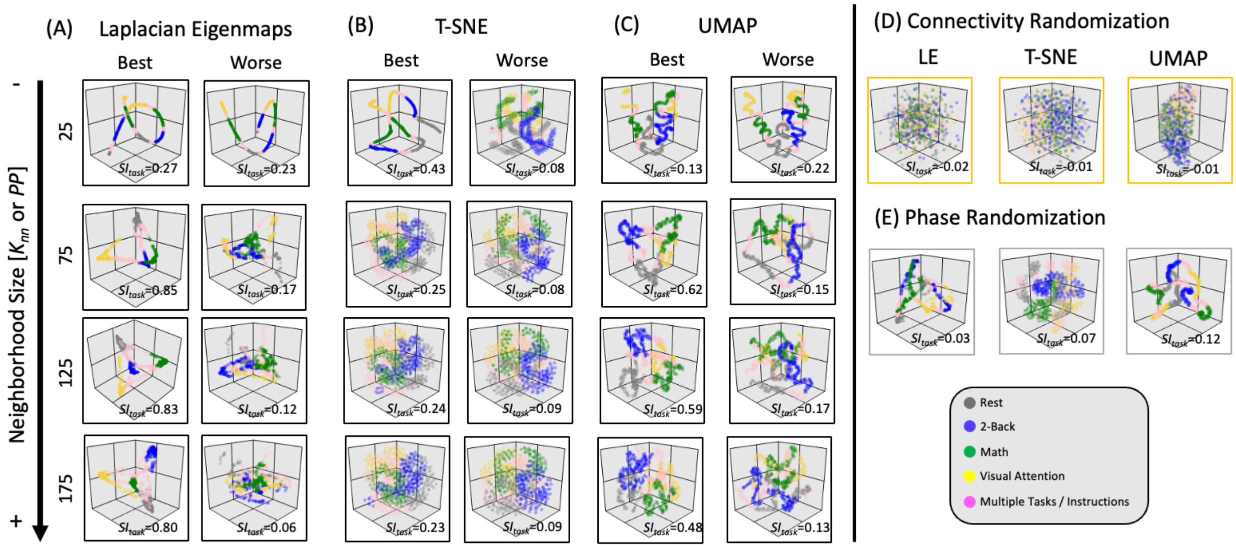


Supplementary Figure 4. Equivalent results to those presented in Figure 7 in the main manuscript, but for 3D instead of 2D.

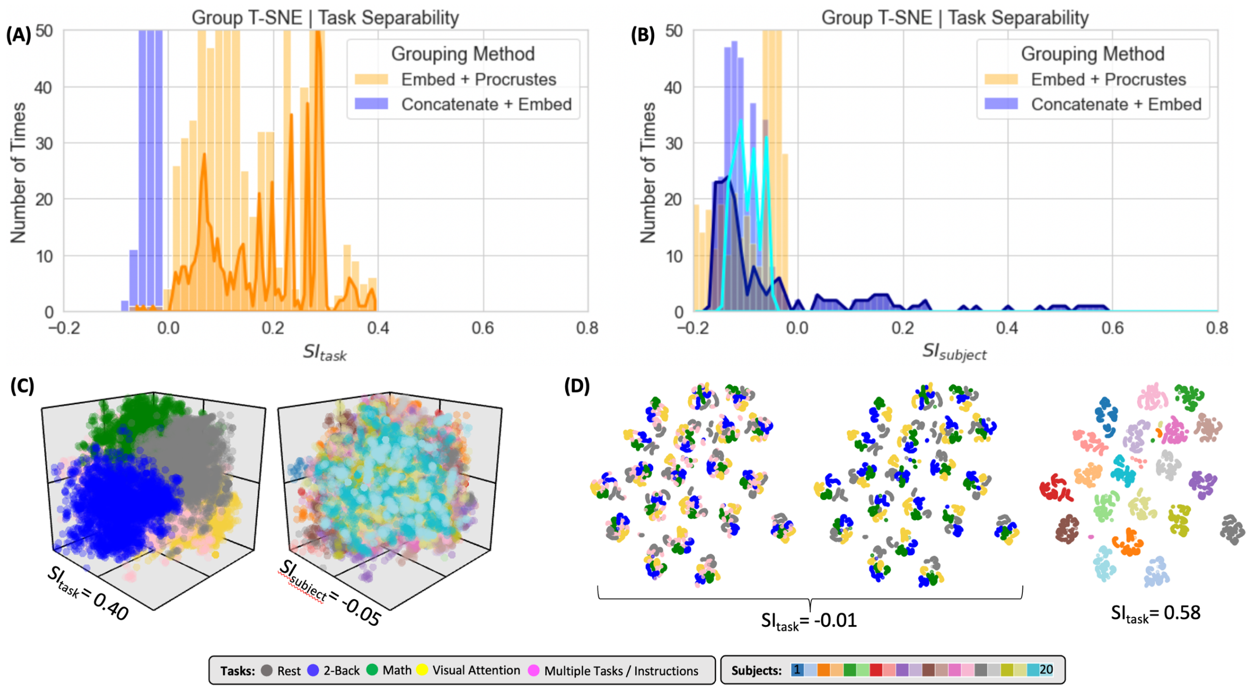


Supplementary Figure 5. Summary of clustering evaluation for T-SNE group embeddings. Due to computational complexity, not all hyper-parameters were explored for group level embeddings and T-SNE. For the "Embed + Procrustes" approach the space explored is restricted to learning rate = 10 or 1000 and the correlation distance. For the "Concatenate + Embed", we only computed embeddings up to m = 10. (A) Histograms of $SI_{task}$ values across all explored hyperparameters when using *Correlation* distance on real data. Distributions are segregated by grouping method: "*Embed + Procrustes*" in orange and "*Concatenation + Embed*" in blue. Dark orange outline highlights the section of the "*Embed + Procrustes*" distribution that corresponds to instances where more than 3 dimensions were used to compute the Procrustes transformation. (B) Histograms of $SI_{subject}$ values across all explored hyperparameters when using *Correlation* distance on real data. Distributions are segregated by grouping method in the same way as in (A). Light blue contour highlights the part of the distribution for "*Concatenation + Embed*" computed on data that has been normalized (e.g., Z-scored), while the dark blue contour highlights the portion corresponding to data that has not been normalized. (C) High quality group-level *"Embed + Procrustes"* embedding annotated by task (left) and subject identity (right). This embedding corresponds to the following hyper-parameter combination: norm = none, PP = 90, dist = correlation, learning rate = 10 and number of dimensions used during Procrutes transformation = 25. (D) High quality group-level "*Concatenation + Embed*" annotated by task (left: including task in-homogenous windows, middle = only task homogenous windows) and subject identity (right). This embedding corresponds to the following hyper-parameter combination: norm = none, PP = 150, dist = correlation, learning rate = 1000 and number of dimensions = 2.

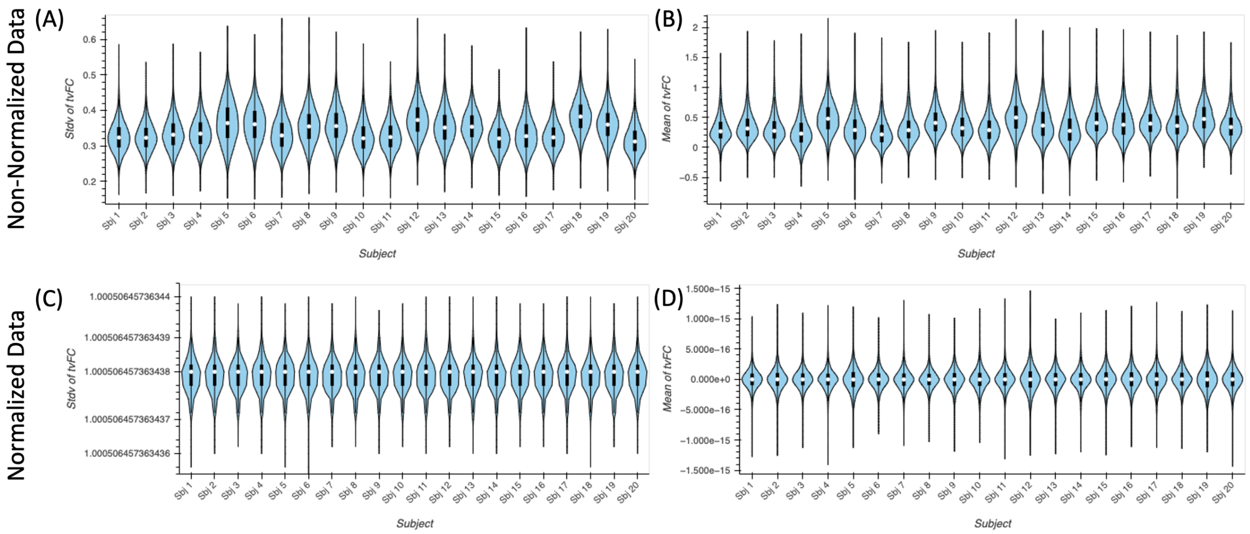


Supplementary Figure 6. (A) Distribution of average FC values across time for all connections separately for each subject prior to data normalization. (B) Distribution of the standard deviation across time of FC values for all connections separately for each subject prior to data normalization. (C) Same as (A) following data normalization. (D) Same as (B) following data normalization.

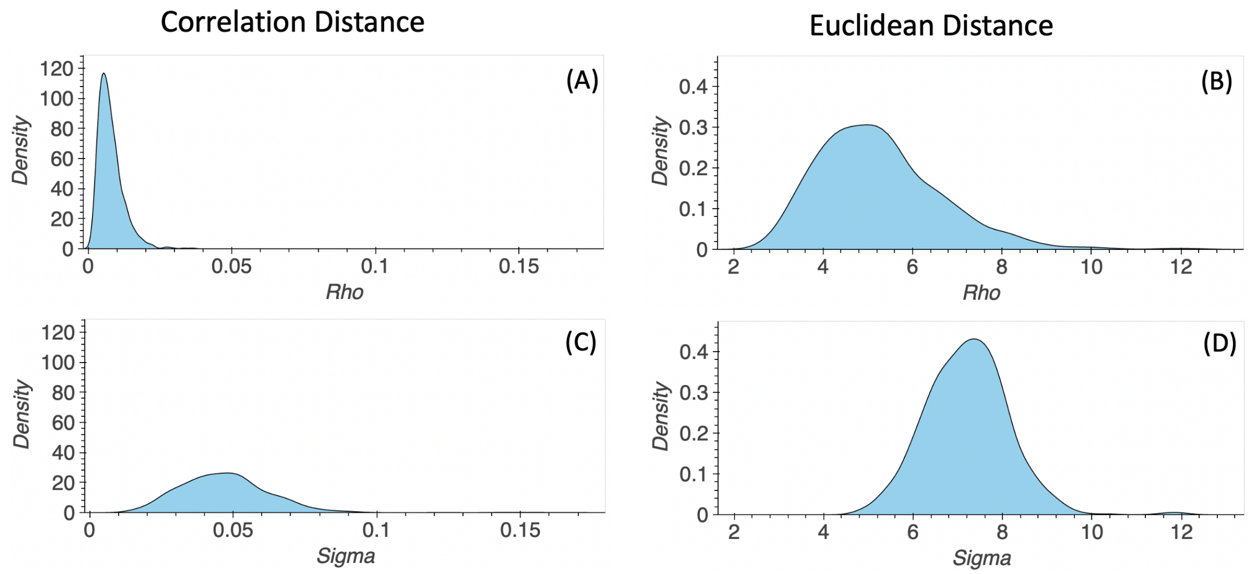


Supplementary Figure 7. Representative distributions of Rho and Sigma values used in the distance normalization step in UMAP when using Correlation (left) and Euclidean distance (right). Results shown here correspond to the following hyper-parameter set ($K_{nn}$ = 90, minimum distance = 0.8, learning rate = 0.01, number of final dimensions = 2). (A) Distribution of Rho values when using the Correlation distance. (B) Distribution of Rho values when using the Euclidean distance. (C) Distribution of Sigma values when using the Correlation distance. (D) Distribution of Sigma values when using the Euclidean Distance.
